# Intersectin and Endophilin condensates prime synaptic vesicles for release site replenishment

**DOI:** 10.1101/2023.08.22.554276

**Authors:** Tyler Ogunmowo, Christian Hoffmann, Renee Pepper, Han Wang, Sindhuja Gowrisankaran, Annie Ho, Sumana Raychaudhuri, Benjamin H. Cooper, Ira Milosevic, Dragomir Milovanovic, Shigeki Watanabe

## Abstract

Neurotransmitter is released from dedicated sites of synaptic vesicle fusion within a synapse. Following fusion, the vacated sites are replenished immediately by new vesicles for subsequent neurotransmission. These replacement vesicles are assumed to be located near release sites and used by chance. Here, we find that replacement vesicles are clustered around this region by Intersectin-1. Specifically, Intersectin-1 forms dynamic molecular condensates with Endophilin A1 near release sites and sequesters vesicles around this region. In the absence of Intersectin-1, vesicles within 20 nm of the plasma membrane are reduced, and consequently, vacated sites cannot be replenished rapidly, leading to depression of synaptic transmission. Similarly, mutations in Intersectin-1 that disrupt Endophilin A1 binding result in similar phenotypes. However, in the absence of Endophilin, this replacement pool of vesicles is available but cannot be accessed, suggesting that Endophilin A1 is needed to mobilize these vesicles. Thus, our work describes a distinct physical region within a synapse where replacement vesicles are harbored for release site replenishment.

## Introduction

At chemical synapses, signaling is mediated by the calcium-dependent exocytosis of synaptic vesicles^1,2^. In resting synapses, synaptic vesicles can dock at release sites within the active zone, where release machinery is concentrated^3–10^. At any given synapse, there exist far fewer release sites than vesicles, and at each release site, only one vesicle can dock^11–13^. This limitation sets an upper boundary for the number of vesicles that are release-ready at any given time. Thus, for a synapse to resist depression of synaptic transmission or to enhance synaptic signaling, the vacated sites must be actively replenished. As such, this replenishment is rate-limiting for continued neurotransmitter release.

Traditionally, the replenishment of release sites was thought to be slow, requiring ∼2-10 s^14–17^. However, synapses are capable of resisting depression of neurotransmitter release during high-frequency stimulation and even enhancing transmission within tens of milliseconds^12,18,19^, suggesting that release site replenishment can be rapid. In line with this, electrophysiological recordings paired with mathematical modeling suggest that there exists a pool of vesicles that respond to docked vesicle depletion and supply new vesicles from so-called ‘replacement sites’ to release sites after fusion^20^. Furthermore, ultrastructural analysis suggests that vesicles can transiently dock to resist depression and enhance synaptic strength in a calcium-dependent fashion^7,21^, a process that may reflect the transition of a replacement vesicle to a docked vesicle. Thus, release sites can be replenished on a millisecond time scale after fusion events^20,22–25^.

There are several functional pools of synaptic vesicles, including the readily-releasable pool, reserve pool, and recycling pool^11,26–28^. Recent studies suggest that some of these pools are separated into distinct physical domains within a synapse by a molecular condensation process^29^. For example, the readily-releasable pool of vesicles may be organized by the condensation of active zone proteins such as RIM, RIM-BP2, voltage-gated Ca^2+^ channels, Munc13-1, ELKS-1 and Liprin-α^30–32^. Additionally, a few synaptic vesicles are thought to tether to these active zone phases^33^. Endocytic zones at presynapses are organized by a specific splice variant of Dynamin 1, Dyn1xA, and its binding partner, Syndapin 1, and these proteins control the endocytic flux of synaptic vesicles^34^. Similarly, the reserve pool of synaptic vesicles is separated by the multivalent interactions of the synaptic vesicle binding protein Synapsin 1 (Syn1) on vesicle membranes^35,36^. Thus, synaptic vesicles are physically organized by molecular condensation, engendering specific functional roles at synapses.

The reserve pool is suggested to also contain proteins like Intersectin-1 (Itsn1) and Endophilin A1 (EndoA1)^37^. Interestingly, Itsn1 and EndoA1 interact with each other and may form condensates on some of these vesicles either independently of or in concert with Syn1^36,38–40^. However, Itsn1 and EndoA1’s function may lie outside of the reserve pool. Non-neuronal secretory cells display Itsn1 enrichment at sites of granule secretion, and its knockdown perturbs hormone release during sustained activity^41^. More recently, work in neuroendocrine adrenal chromaffin cells shows that Itsn1 together with EndoA1 mediates granule replenishment during stimulation^42^. At the calyx of Held, fast vesicle replenishment seems to be abolished in Itsn1 knockout (KO) neurons^43^. Furthermore, absence of either Itsn1 or EndoA1 in mouse hippocampal synapses leads to synaptic depression during a train stimulus^44,45^x,. These data suggest that Itsn1 and EndoA1 may contribute to activity-dependent replenishment of release sites.

Here, we demonstrate that Itsn1-EndoA1 form condensates and mediate the maintenance of the replacement vesicle pool. Specifically, Itsn1 and EndoA1 are located between the active zone and the Syn1-positive reserve vesicle cluster. In neurons lacking Itsn1 or the functional interaction of Itsn1 with EndoA1, the number of vesicles within 20 nm of the active zone is significantly reduced. Without EndoA1, replacement vesicles are intact, yet not accessible for docking. Consequently, the replenishment of release sites by transient docking is slowed, leading to accelerated depression of synaptic transmission. These data suggest that a subset of replacement synaptic vesicles are held near the active zone and used in an activity-dependent manner to support neurotransmission during repetitive stimulation.

## Results

### Intersectin-1 forms molecular condensates with synaptic proteins and vesicles

Intersectin-1 Long (L) (hereafter, Itsn1) is a neuronally enriched isoform of Itsn1 (Fig. 1a). Itsn1 has five Src-Homology 3 (SH3) domains in tandem (namely, A-E; Fig. 1a), which enable interaction with a multitude of synaptic proteins, including Synapsin 1 (Syn1) and Endophilin A1 (EndoA1)^35,36,38,40,46^. When EGFP (hereafter, GFP)-Itsn1 was overexpressed alone in HEK293T cells, Itsn1 readily formed condensed molecular structures (Fig. 1b), likely due to the interactions with the endogenously expressed proteins with proline-rich motifs. These molecular condensates spontaneously fused with one another (Fig. 1c), as is typical with molecular condensates^47^. These Itsn1 condensates were dispersed by the application of 1,6-Hexanediol (Fig. 1d), an aliphatic alcohol that disrupts weak multivalent interactions^48^. Further, internal fluorescence recovery after photobleaching (FRAP) measurements revealed that fluorescence recovery within Itsn1 condensates depends on the diameter of photobleaching (Fig. 1e-g), suggesting that molecules remain highly mobile within condensates^48^. These data are consistent with the idea that Itsn1 dynamically condenses in cells.

**Fig. 1.**
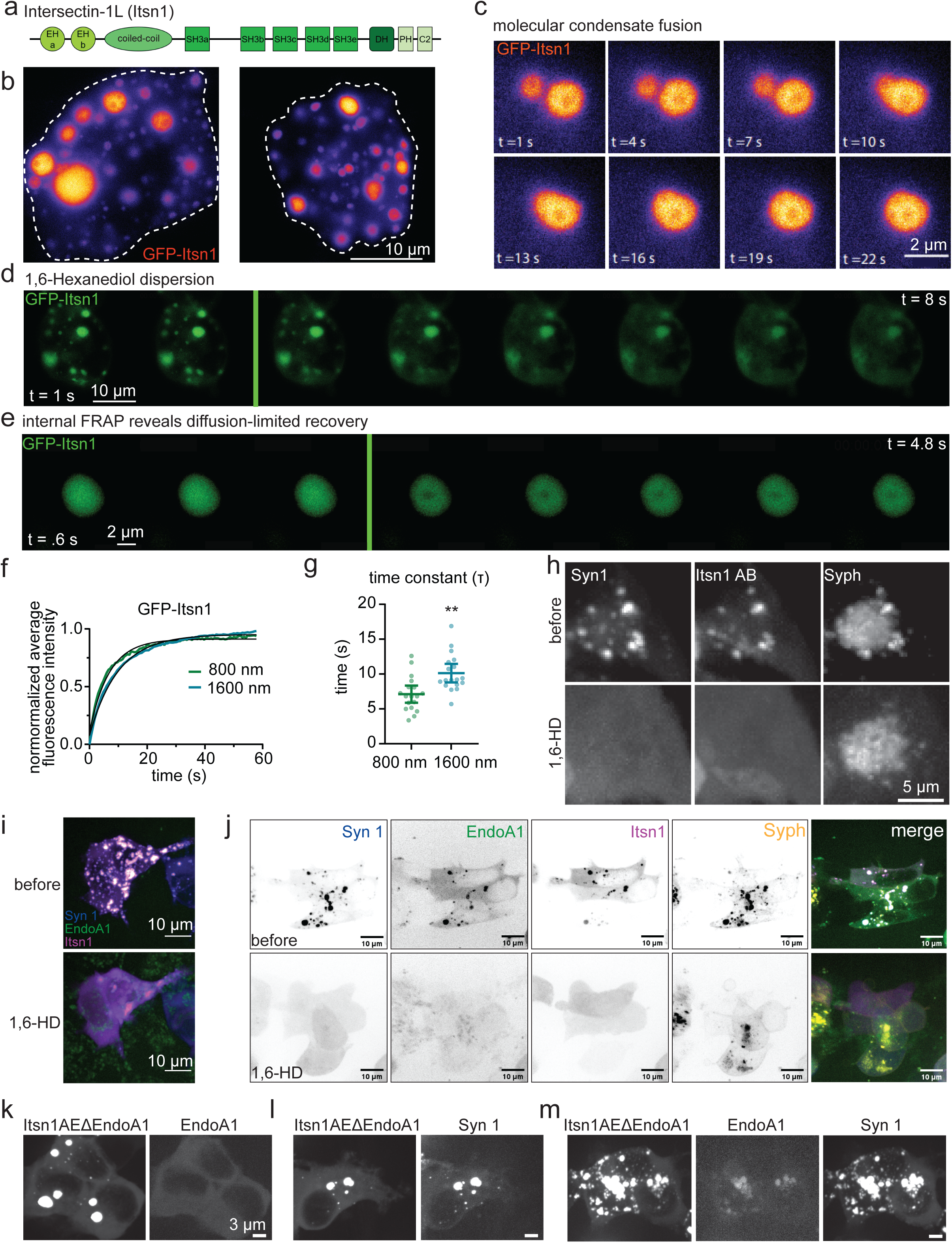
Intersectin 1 forms the condensates with Endophilin A1. a. The protein domain structure of Intersectin-1L (Itsn1). b. Two HEK293T cells expressing GFP-Itsn1 full-length (FL). Scale bar, 10 µm. c. Example live-HEK293T cell images showing GFP-Itsn1 FL condensates undergoing a fusion event. Times after the initiation of image acquisitions are indicated. Scale bar, 2 µm. d. Example live-cell fluorescence micrographs showing GFP-Itsn1 signals over 8 s with the addition of 4% 1,6-Hexanediol at 3 s. Scale bar, 10 µm. e. Fluorescence recovery of GFP-Itsn1 signals when signals within the condensate was photobleached with a diameter of 800 or 1600 nm. Each frame represents 0.6 s. Scale bar, 2 µm. f. Plot showing fluorescence recovery after photobleaching with a bleaching spot of the indicated diameters in HEK293T cells expressing GFP-Itsn1. g. Plot showing time constant Tau of fluorescence recovery in f. Bars are the mean; error bars are SEM. Student’s t test. p**<0.01. h. (top) HEK293 cells co-expressing mCherry-Synapsin 1 (Syn1), GFP-Itsn1 AB, and Synaptophysin-emiRFP670 (Syph). (bottom) Cells in (top) after 3% 1,6 Hexanediol treatment. Scale bar, 5 µm. i. (top) HEK293 cells co-expressing BFP-Syn1, mCerulean-Endophilin A1 (EndoA1), and GFP-Itsn1 AB. (bottom) cells in (top) after 3% 1,6 Hexanediol treatment. Scale bars, 10 µm. j. (top) HEK293 cells co-expressing BFP-Syn1, mCerulean-EndoA1, GFP-Itsn1 AB, and Syph-emiRFP670. Each individual channel is separated, and then merged. (bottom) cells in (top) after 4% 1,6 Hexanediol treatment. Scale bars, 10 µm. k-m. HEK293 cells co-expressing mutant mCherry-Itsn1 (Itsn1 W949E and Y965E; Itsn1AEΔEndoA1) and mCerulean-EndoA1 (l), mCherry-Itsn1 EΔEndoA1 and BFP-Syn1 (m), and mCherry-Itsn1AEΔEndoA1, mCerulean-EndoA1, and BFP-Syn1 (n). Scale bar, 5 µm. See Supplementary Table 1 for additional information.

Recently, an SH3 A-E concatemer of Itsn1 is shown to regulate the assembly of Syn1 condensates in a biphasic fashion and in a concentration-dependent manner^35^, suggesting Itsn1 condensates may interact with and modulate other presynaptic phases. To assess whether Itsn1 can modulate the boundary properties of Syn1 condensates, we used GFP-tagged concatemers of Itsn1 containing either only two (Itsn1 AB) or five (Itsn1 AE) SH3 domains in HEK293 cells (Extended Data Fig. 1)^35^. Consistent with previous work^35^, co-expression of mCherry-Syn1 with these GFP-tagged concatemers of Itsn1 readily led to the formation of molecular condensates (Extended Data Fig. 1a). In line with the fluid-nature of Synapsin condensates^35^, an increase in the valency of SH3 concatemers (i.e., two vs. five) and the expression time (i.e., protein concentration in the cytosol) causes these condensates to become larger and less numerous over time (Extended Data Fig. 1a,b). Additionally, these condensates were dispersed by the application of 1,6-Hexanediol (Extended Data Fig. 1c,d)^47^. These data suggest that Itsn1 regulates the material properties of Syn1 condensates.

Previous studies show that the expression of Syn1 and Synaptophysin 1 (Syph) in non-neuronal cells leads to the formation and clustering of synaptic vesicle-like organelles^39^. To investigate the role of Itsn1 in regulating the Syn1-mediated clustering of synaptic vesicles, we expressed GFP-Itsn1 AB with mCherry-Syn1 and Syph-emi-RFP670. Itsn1 colocalized with Syn1 and Syph (Fig. 1h), suggesting that these proteins co-assemble on these synaptic vesicle-like organelles. These vesicle-containing condensates were also sensitive to the 1,6-Hexanediol treatment (Fig. 1h), indicating their fluid-like property is not changed by the presence of these synaptic vesicle-like organelles. In line with this, expression of untagged Syph did not change the size or number of Itsn1-Syn1 condensates (Extended Data Fig. 1e). These findings suggest that Itsn1-Syn1 condensates can contain these synaptic vesicle-like organelles without affecting their co-assembly.

### EndoA1 coalesces with Itsn1 condensates

Recently, EndoA1 was shown to both facilitate the phase separation of Syn1 and enter synaptic vesicle-like clusters through interaction with Syn1^40^. Further, this Syn1-EndoA1 phase contains a variety of synaptic proteins including Itsn1. To test whether EndoA1 is also found in Itsn1 condensates, we expressed mCerulean-EndoA1 with GFP-Itsn1 AB or GFP-Itsn1 AE. We also expressed BFP-Syn1 in place of GFP-Itsn1 AE as a positive control, and EndoA1 alone. Unlike GFP-Itsn1, mCerulean-EndoA1 expressed alone was diffuse within cells (Extended Data Fig. 1f). However, as in the previous study^40^, EndoA1 formed 1,6-Hexanediol-sensitive condensates with Syn1 when co-expressed (Extended Data Fig. 1g). When co-expressing GFP-Itsn1 AB, mCerulean-EndoA1, and BFP-Syn1, 1,6-Hexanediol sensitive condensates also formed in cells (Fig. 1i). Importantly, Itsn1 condensates readily incorporated EndoA1 in the absence of Syn1 (Extended Data Fig. 1h), suggesting these proteins can assemble into condensates on their own. Condensates formed by differential combinations of Itsn1 AB, Itsn1 AE, Syn1 and EndoA1 all displayed distinct circularities and sizes (Extended Data Fig. 1i,j), yet they formed within cells at the same frequency (Extended Data Fig. 1k), suggesting molecular composition changes condensate properties but does not affect their formation likelihood. When co-expressed with Syph-emiRFP670, Itsn1-EndoA1 condensates formed, and they were dispersed by 1,6-Hexanediol (Extended Data Fig. 1l). However, unlike Itsn1-Syn1 condensates, they did not contain Syph-emiRFP670 signal (Extended Data Fig. 1l), indicating that some level of Syn1 is required to initiate clustering within Itsn1 condensates. In fact, when we expressed all four proteins (GFP-Itsn1 AB, mCerulean-EndoA1, BFP-Syn1, and Syph-emiRFP670), they formed condensates (Fig. 1j) that can be dispersed by 1,6-Hexanediol (Fig. 1j), suggesting that these proteins together can cluster vesicles. Lastly, this coalescence of EndoA1 with Itsn1 required their specific interaction, an unconventional SH3-SH3 interaction^38^. When Itsn1 AE’s SH3B domain (W949E/Y965E) was mutated to block EndoA1 interaction (Itsn1 AE ΔEndoA1)^38^, Itsn1 condensates completely lacked EndoA1 (Fig. 1k). This is likely due to the lack of direct biochemical interaction - Itsn1ΔEndoA1 condensates displayed FRAP recovery kinetics identical to wild-type (WT) GFP-Itsn1 (Extended Data Fig. 1m,n), suggesting the physical properties of these condensates are not changed by these mutations. Consistently, Itsn1 AE ΔEndoA1 condensates incorporated Syn1 normally (Fig. 1l). Interestingly, if Syn1 and Itsn1 AE ΔEndoA1 are co-transfected with EndoA1, EndoA1 is found within Itsn1-Syn1 condensates (Fig. 1m), indicating that Syn1 condensates facilitate the accumulation of both Itsn1 and EndoA1 regardless of their direct interaction, likely through Syn1’s own interactions with both proteins^36^. Together, our results suggest that Itsn1 can form dynamic assemblies on vesicles in the presence of Syn1, which can contain EndoA1, indicating that Itsn1 condensates can potentially regulate vesicle dynamics in concert with EndoA1 and Syn1.

### Intersectin-1 and Endophilin A1 colocalize on synaptic vesicles near active zones

To test whether Itsn1 and EndoA1 form condensates in synapses, we first performed 2D stimulated emission depletion (STED) microscopy and Instant Structured Illumination Microscopy (ISIM) to localize these proteins (Fig. 2a-g). We visualized the relative location of Itsn1 and EndoA1 to the reserve pool marked by Syn1, the active zone marked by Bassoon or RIM, and synaptic vesicles more broadly marked by Synaptobrevin-2 (Syb2). In 2D STED, mouse hippocampal neurons expressing Itsn1 or EndoA1 tagged with GFP were stained with an anti-GFP antibody together with either Syb2 or Bassoon antibodies. Fluorescent puncta were quantified as a function of their distance from the boundary of either Bassoon or Syb2 signals^34^. Both Itsn1 and EndoA1 were localized primarily adjacent to the Bassoon boundary (active zone), with ∼20% located within the active zone boundary and ∼80% located outside, highly enriched right at the boundary line. (Fig. 2a and Extended Data Fig. 2a). When co-stained with Syb2, Itsn1 and EndoA1 signals peaked on the boundary of Syb2 signals (synaptic vesicle-enriched regions), with ∼40% located within the Syb2 boundary and ∼60% located outside (Fig. 2b and Extended Data Fig. 2b). The average distance of Itsn1 and EndoA1 puncta to Bassoon or Syb2 boundaries was identical, with puncta being on average ∼2 times closer to Syb2 (Fig. 2c). These data suggest that a significant fraction of Itsn1 and EndoA1 proteins are contained within and next to the active zone, and an even larger fraction is on or near synaptic vesicles.

**Fig. 2.**
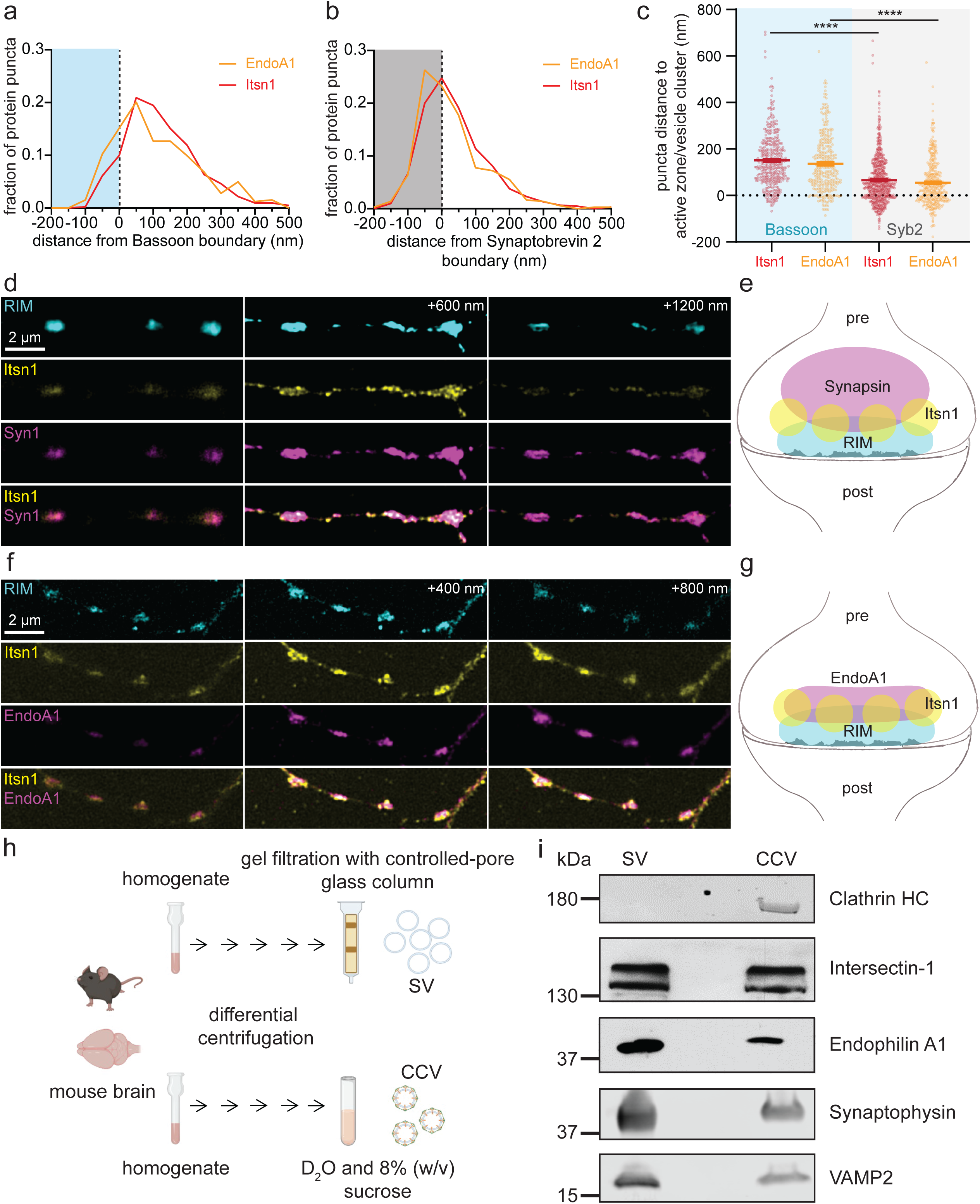
Itsn1 and EndoA1 colocalize near the active zone. a. Cumulative plots showing the distribution of GFP-Itsn1 and GFP-EndoA1 puncta by 2D STED relative to the active zone boundary (dotted line), defined by Bassoon signal (blue fill). b. Cumulative plots showing the distribution of GFP-Itsn1 and GFP-EndoA1 puncta by 2D STED relative to synaptic vesicle localization boundary (dotted line), defined by Syb2 signal (gray). c. Plot showing distances between either GFP-Itsn1 or EndoA1-GFP puncta and either the active zone boundary or synaptic vesicle localization boundary. Bars are the mean; error bars are SEM. Kruskal-Wallis test, with Dunn’s multiple comparisons test. ****p<0.0001. p values are shown for comparisons between both GFP-Itsn1 and EndoA1-GFP distances to Syb2 versus distances to Bassoon. d. Example presynapses visualized by ISIM. En face view shown. Endogenous RIM was stained by an anti-RIM antibody and a secondary antibody conjugated to Alexa488. Endogenous Itsn1 was stained by an anti-Itsn1 antibody and a secondary antibody conjugated to Alexa568. Endogenous Synapsin was stained by an anti-Synapsin antibody and a secondary antibody conjugated to Alexa647. Each z-slice shown is separated by 600 nm. Scale bar, 2 µm. e. Schematic of the relative Synapsin, Itsn1 and RIM localization within presynapses (pre). f. Example presynapses visualized by ISIM. En face view shown. Endogenous Rim was stained as in d. Endogenous Itsn1 was stained as in d. Endogenous EndoA1 was stained by an anti-EndoA1 antibody and a secondary antibody conjugated to Alexa647. Each z-slice is separated by 400 nm. Scale bar, 2 µm. g. Schematic of the relative EndoA1, Itsn1, and RIM localization within presynapses (pre). Itsn1 and EndoA1 signals largely overlap. h. Pipeline schematic for synaptic vesicle (SV) and clathrin-coated vesicle (CCV) isolation. i. Blots showing the amount of indicated proteins on purified synaptic vesicles (SV) and clathrin-coated vesicles (CCV). See Supplementary Table 1 for additional information.

For the 3D organization of Itsn1 and EndoA1 proteins, we performed ISIM imaging, using both the active zone marked by RIM and the reserve pool marked by Syn1. Here, Itsn1 and EndoA1 were endogenously stained by antibodies. ISIM revealed that Itsn1 forms small puncta at synapses, and these puncta appeared to be sandwiched between RIM signals and Syn1 signals (Fig. 2d,e and Extended Data Fig. 2c). Itsn1 colocalized with EndoA1 just above the RIM signals (Fig. 2f,g and Extended Data Fig. 2d). Additionally, some Itsn1 puncta were also found outside the synapses (Fig. 2d and Extended Data Fig. 2c,d). These data suggest that Itsn1 and EndoA1 form puncta near the active zone.

To test whether these puncta represent molecular condensates formed on vesicles, we first applied 7% 1,6-Hexanediol to neurons expressing GFP-Itsn1 and mCherry-Syn1 (Extended Data Fig. 3) and measured the coefficient of variation (CV) of fluorescent signals along the axons to quantify condensate dispersion^40^. Within 30 s, puncta in the soma (Extended Data Fig. 3a-c) and the axons (Extended Data Fig. 3d-e) of these neurons were markedly diffused. In agreement, measured CVs for both axonal GFP-Itsn1 and axonal mCherry-Syn1 signals were reduced after 1,6-Hexanediol treatment (Extended Data Fig. 3f-g), indicating that condensed structures are dispersed and thus, are formed via the weak hydrophobic interactions of these proteins. We then tested whether these condensates undergo dynamic dispersion and reclustering during neuronal activity by expressing either GFP-Itsn1 or EndoA1-mRFP in WT neurons and following its distribution after 300 action potentials (APs) given at 10 Hz (Extended Data Fig. 3h-j). As in previous reports, which suggest Endophilin disperses from synapses during activity^40^, these two proteins undergo dynamic dispersion and then recondense in response to neuronal activity (Extended Data Fig. 3i,j). Finally, to test if these proteins condense onto synaptic vesicles, we purified synaptic vesicles from mouse brains^49^ and probed for Itsn1 and EndoA1 using antibodies (Fig. 2h). We used purified clathrin-coated vesicles as a control for staining since Itsn1 and EndoA1 are found on these vesicles^49^. We found both Itsn1 and EndoA1 are present on purified synaptic vesicles (Fig. 2h,i). Together, these data suggest that Itsn1 and EndoA1 form condensates in synapses and are likely localized on synaptic vesicles that are present near the active zone^39^.

### Intersectin-1 maintains a vesicle pool for transient docking

Our data so far suggests that Itsn1 and EndoA1 form condensates on synaptic vesicles near the active zone. To discern the synaptic function of these condensates, we conducted zap-and-freeze time-resolved electron microscopy experiments^7^, which visualize synaptic membrane trafficking with millisecond temporal precision by electron microscopy. *Itsn1+/+* (WT) and *Itsn1-/-* (KO) mouse hippocampal neurons were frozen either unstimulated or stimulated with a 1-ms electrical pulse, which induces a single AP, at various time points prior to freezing (Fig. 3 and Extended Data Fig. 4)^7^. Since Itsn1 was previously implicated in endocytosis in mouse hippocampal synapses^50^ and in the *Drosophila* neuromuscular junction^51^, we first quantified endocytic pit formation and resolution by stimulating Itsn1 WT or KO neurons and freezing 100 ms or 1 s after^5^. Consistent with recent work^45^, we found no significant defect in ultrafast endocytosis in Itsn1 KO synapses (Fig. 3c), suggesting that Itsn1 is not essential for synaptic vesicle recycling in these synapses.

**Fig. 3.**
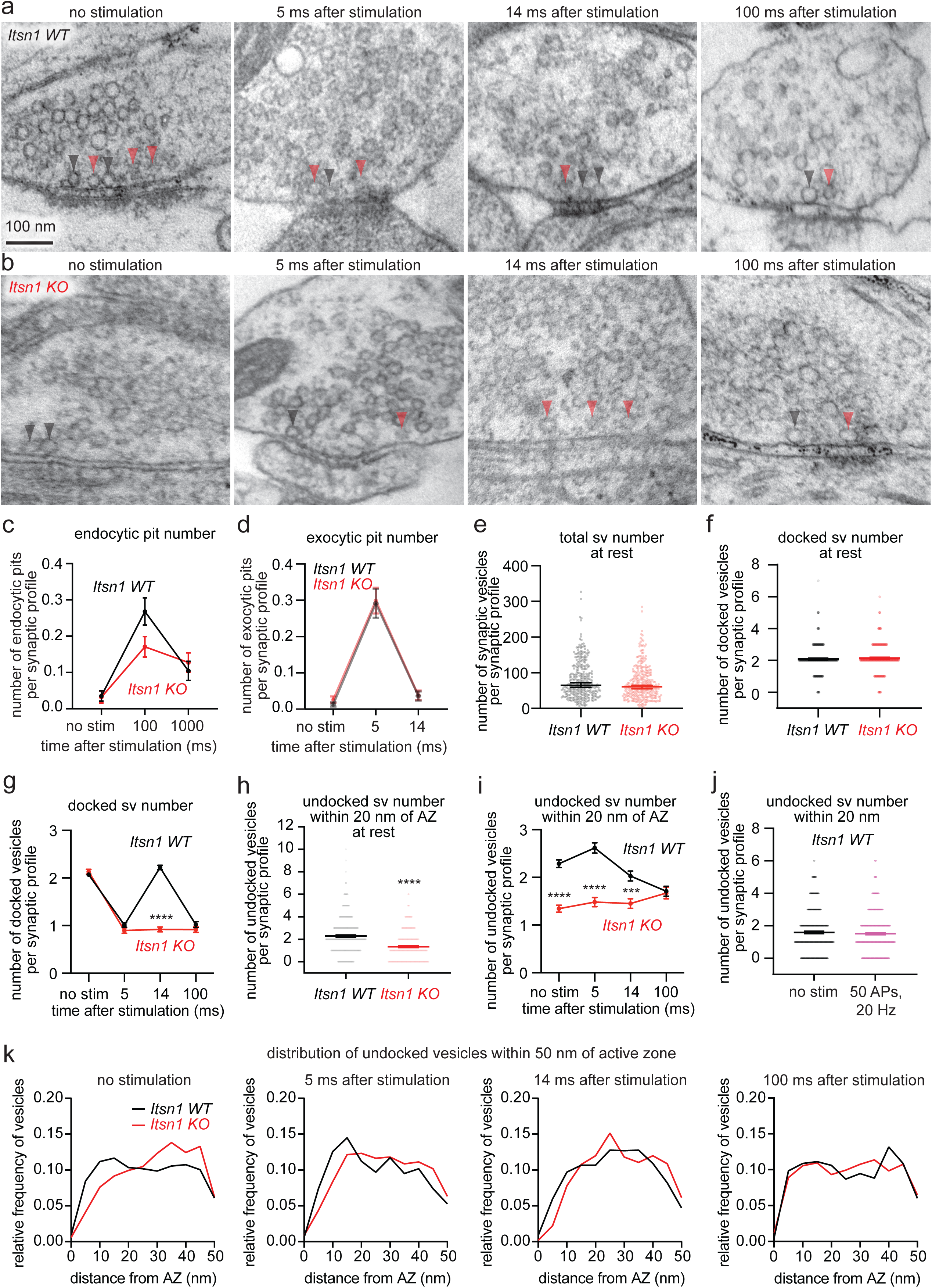
Intersectin 1 organizes undocked vesicles needed for transient docking. a. Electron micrographs showing the progression of docked and undocked vesicle abundance and localization at rest and at indicated time points following stimulation (1 ms electric pulse, 37 °C, 4 mM external Ca^2+^) in *Itsn1+/+* (WT). Black arrowhead: docked vesicle. Red arrowhead: undocked vesicle. Docked vesicles are defined as those in physical contact with plasma membrane. Scale bar, 100 nm. b. Electron micrographs showing vesicle dynamics as described in a. for *Itsn1-/-* (KO) synaptic profiles. c. Plots showing the number of endocytic pits in *Itsn1 WT* (black) and *KO* (red) synaptic profiles at rest, 100 ms, and 1000 ms after stimulation. Dots are the mean; error bars are SEM. d. Plots showing the number of exocytic pits in *Itsn1 WT* (black) and *KO* (red) synaptic profiles at rest, 5 ms, and 14 ms after stimulation. Dots are the mean; error bars are SEM. e. Number of synaptic vesicles per synaptic profile in *Itsn1 WT* (black) and *KO* (red) synaptic profiles. Bars are the mean; error bars are SEM. f. The total number of vesicles docked at the active zone at rest in *Itsn1 WT* and *Itsn1 KO* synaptic profiles. Bars are the mean; error bars are SEM. g. Number of docked vesicle at rest and 5, 14, and 100 ms after stimulation in *Itsn1 WT* and *Itsn1 KO* synaptic profiles. Dots are the mean; error bars are SEM. Kruskal-Wallis test, with Dunn’s multiple comparisons test. ****p<0.0001. Comparisons were made between *Itsn1 WT* and *Itsn1 KO*. h. Number of undocked vesicles within 20 nm of the active zone membrane at rest in *Itsn1 WT* and *Itsn1 KO* synaptic profiles. Bars are the mean; error bars are SEM. Mann-Whitney U test. ****p<0.0001. i. Number of undocked vesicles within 20 nm of the active zone at rest and 5, 14, and 100 ms after stimulation in *Itsn1 WT* and *Itsn1 KO* synaptic profiles. Bars are the mean; error bars are SEM. Kruskal-Wallis test, with Dunn’s multiple comparisons test. ****p<0.0001, ***p<0.001. Comparisons were made between *Itsn1 WT* and *Itsn1 KO*. j. Number of undocked vesicles within 20 nm of the active zone in wild-type synaptic profiles at rest or after 50 repetitive stimuli delivered at 20 Hz. k. Relative frequency distributions of undocked vesicles 2-50 nm from the active zone membrane. Vesicle counts were determined in *Itsn1 WT* and *Itsn1 KO* synaptic profiles at rest, 5, 14, and 100 ms after stimulation (left to right). Vesicle counts were separated into 2 nm bins and normalized by the total number of vesicles in this region. AZ = active zone. See Supplementary Table 1 for additional information.

To further probe the role of Itsn1 at synapses, we assessed vesicle dynamics at synapses. We defined vesicles touching the plasma membrane as docked, as in our previous studies^4,5,7^. In unstimulated conditions, Itsn1 KO neurons had a normal number of docked and undocked vesicles (Fig 3a,b,e,f, and Extended Data Fig. 4a,b). To test whether these synapses functioned normally, we stimulated these neurons once and froze 5 ms after. Both Itsn1 WT and KO displayed exocytic pits at this time point (Fig. 3d). The number of exocytic pits in both cases is nearly identical, indicating no major exocytic phenotypes (Fig. 3d). Concomitantly, the number of docked vesicles was reduced normally at 5 ms (Fig. 3a,b,g, Extended Data Fig 4a,b). Thus, exocytosis after a single stimulus is unaffected by Itsn1 KO, consistent with previous studies in the Calyx of Held synapses^43^.

We next observed docking dynamics after stimulation to test whether Itsn1 is involved in refilling release sites. Our previous studies suggest that vacated release sites can be replenished by ∼14 ms after a single stimulus^7^, a process that may reflect synaptic plasticity and maintenance^20,21,24^. This docking process is reversible (hence, ‘transient docking’) - by 100 ms, the number of docked vesicles returns to the depleted state^4,5,7^. To test this functional paradigm in Itsn1 KO synapses, we froze neurons 14 ms and 100 ms after a single stimulus. In Itsn1 WT synapses, transient docking was normal—docked vesicles were recovered at 14 ms and depleted again by 100 ms (Fig. 3a,g, Extended Data Fig. 4a). By contrast, in Itsn1 KO synapses, transient docking was completely abolished (Fig. 3b,g, Extended Data Fig. 4b). Interestingly, undocked vesicles in Itsn1 KO synapses appeared scarce near the active zone, particularly in the first 20 nm (Fig. 3h). This ∼35% reduction was present at baseline and persisted out to 14 ms after stimulation (Fig. 3h,i). Additionally, the distribution of undocked vesicles close to the active zone (within 50 nm) changed dynamically following an action potential. In Itsn1 WT, the number of undocked vesicles within 20 nm transiently increased at 5 ms after stimulation from baseline levels while slightly decreasing within 20-50 nm (Fig. 3i,k). This accumulation of undocked vesicles within 20 nm of the active zone then decreased in number at 14 ms, even slightly below the baseline, likely due to the transition to a docked state mirroring transient docking (Fig. 3i,k). In resting Itsn1 KO synapses, undocked vesicles were more abundant between 20-50 nm of active zones (Fig. 3k), and redistribution of these vesicles into the 20 nm region was slow, requiring ∼100 ms for Itsn1 KO synapses to reach Itsn1 WT levels (Fig. 3i,k). With repetitive stimulation (50 APs, 20 Hz) in WT neurons, vesicles within 20 nm of the active zone were maintained, indicating that during steady turnover of vesicles from this zone to release sites, this zone is actively refilled (Fig. 3j). Taken together, these data suggest that a sufficient number of undocked vesicles must be within 20 nm of the active zone to be used for transient docking. Without Itsn1, these vesicles seem to be located further away from the active zone, preventing their use. Thus, hereafter, we will refer to these vesicles as replacement vesicles, and this 20 nm region as the replacement zone.

### Intersectin-1 and Endophilin A1 interaction is critical for vesicle replacement

Itsn1 and EndoA1 form condensates on vesicles and co-accumulate near the active zone, and without Itsn1, the consequent reduction in replacement vesicles coincides with failure of transient docking. Recent work suggests that the interaction of Itsn1 and EndoA1 is critical for continued vesicle release in stimulated chromaffin cells^42^, highlighting the potential for neuronal vesicle replacement to have similar requirements. To test this possibility, we performed rescue experiments in Itsn1 KO neurons with WT and mutant forms of Itsn1. Itsn1 and EndoA1 interact with one another through an SH3-SH3 interaction^38^. Two point mutations in full length (FL) Itsn1’s SH3B domain (W949E/Y965E^38^; hereafter Itsn1ΔEndoA1) perturbed this interaction (Extended Data Fig. 5a).

At the ultrastructural level, overall synapse morphology was normal, and the total number of vesicles was unperturbed relative to Itsn1 KO when rescue constructs were expressed (Fig. 4a,b, and Extended Data Fig. 5b-d). After a single AP, exocytosis occurred normally^7^ in rescue backgrounds (Extended Data Fig. 5e). When Itsn1 WT was expressed in Itsn1 KO neurons, all phenotypes were rescued (Fig. 4a, c-g, Extended Data Fig. 5b). By contrast, the expression of Itsn1ΔEndoA1 failed to rescue the transient docking phenotype— docked vesicles were depleted at 5 ms normally but failed to return to baseline 14 ms after stimulation (Fig. 4b,c,d, and Extended Data Fig. 5c). As in Itsn1 KO, this failure coincided with a lack of replacement vesicles (Fig. 4b,e,f, and Extended Data Fig. 5c), and the distribution of undocked vesicles was shifted away from the active zone relative to Itsn1 WT rescue in unstimulated synapses, 5 ms and 14 ms after stimulation (Fig. 5g). Taken together, our results suggest that the specific interaction of Itsn1 and EndoA1 is required for the localization of replacement vesicles and their mobilization for transient docking.

**Fig. 4.**
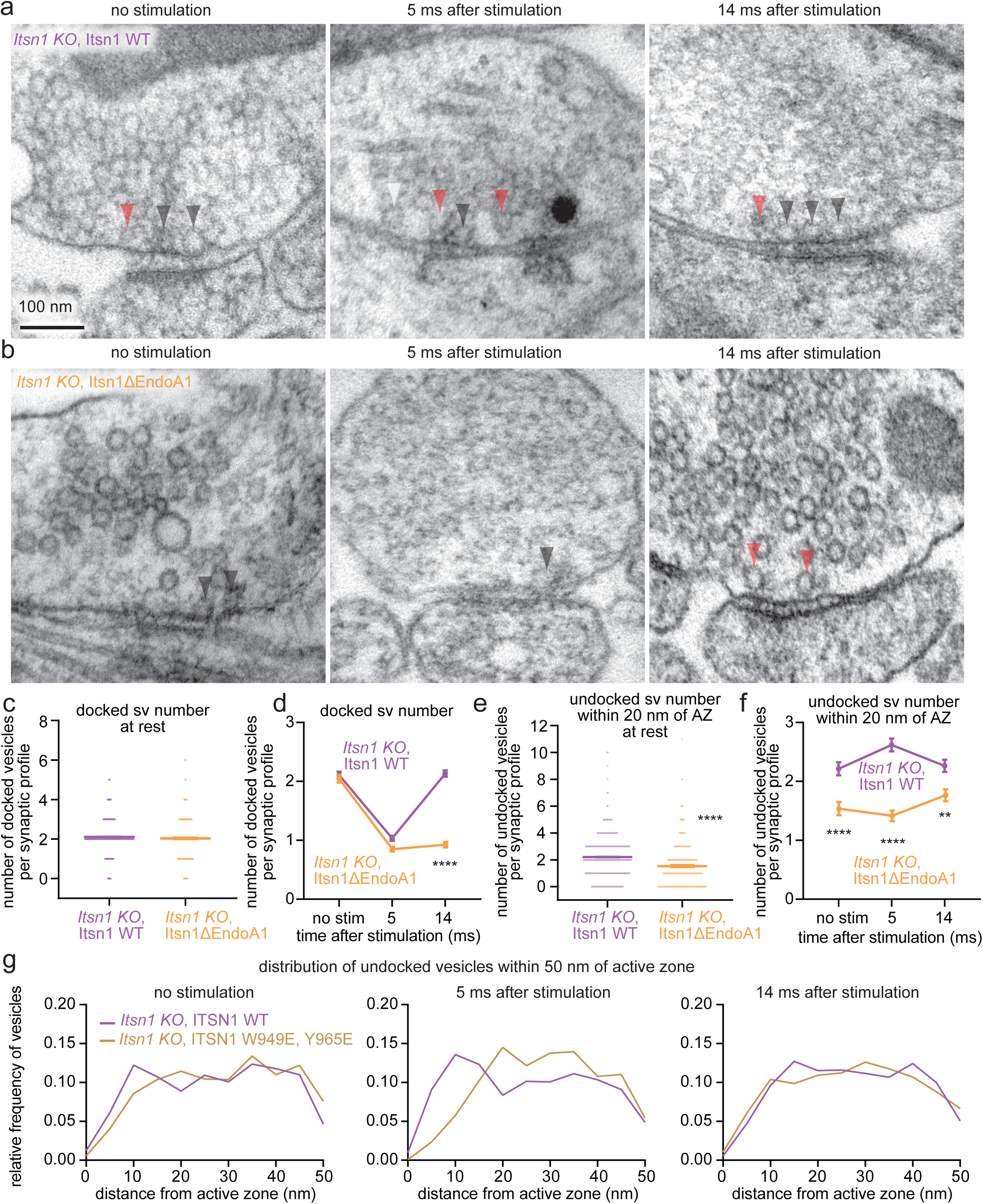
Interaction of Itsn1 and EndoA1 is required for transient docking. a-b. Electron micrographs showing the progression of docked vesicle and undocked vesicle abundance and localization in *Itsn1 KO* neurons expressing Itsn1 WT (a) and Itsn1 W949E, Y965E mutant (Itsn1ΔEndoA1) (b) at rest, 5, or 14 ms after stimulation (1 ms electric pulse, 37 °C, 4 mM external Ca^2+^). Black arrowhead: docked vesicle. Red arrowhead: undocked vesicle. Scale bar, 100 nm. c. Number of docked vesicles in *Itsn1 KO,* Itsn1 WT and *Itsn1 KO*, Itsn1ΔEndoA1 synaptic profiles at rest. Bars are the mean; error bars are SEM. Mann Whitney U test. ****p<0.0001. d. Number of docked vesicles at rest, 5 and 14 ms after stimulation in *Itsn1 KO,* Itsn1 WT and *Itsn1 KO*, Itsn1ΔEndoA1 synaptic profiles. Bars are the mean; error bars are SEM. Kruskal-Wallis test, with Dunn’s multiple comparisons test. ****p<0.0001. Comparisons were made between *Itsn1 KO,* Itsn1 WT and *Itsn1 KO*, Itsn1ΔEndoA1. e. Same as in c, but for undocked vesicles within 20 nm of the active zone. Bars are the mean; error bars are SEM. Mann-Whitney U test. ****p<0.0001. f. Same as in d, but for undocked vesicles within 20 nm of the active zone. Bars are the mean; error bars are SEM. Kruskal-Wallis test, with Dunn’s multiple comparisons test. **p<0.01, ****p<0.01. Comparisons were made between *Itsn1 KO,* Itsn1 WT and *Itsn1 KO*, Itsn1ΔEndoA1. g. Relative frequency distributions of undocked vesicles 2-50 nm from the active zone membrane in *Itsn1 KO,* Itsn1 WT and *Itsn1* KO, Itsn1ΔEndoA1 synaptic profiles at rest, 5, and 14 ms after stimulation (left to right). Vesicle counts were separated into 2 nm bins and normalized by the total number of vesicles in this region. See Supplementary Table 1 for additional information.

**Fig. 5.**
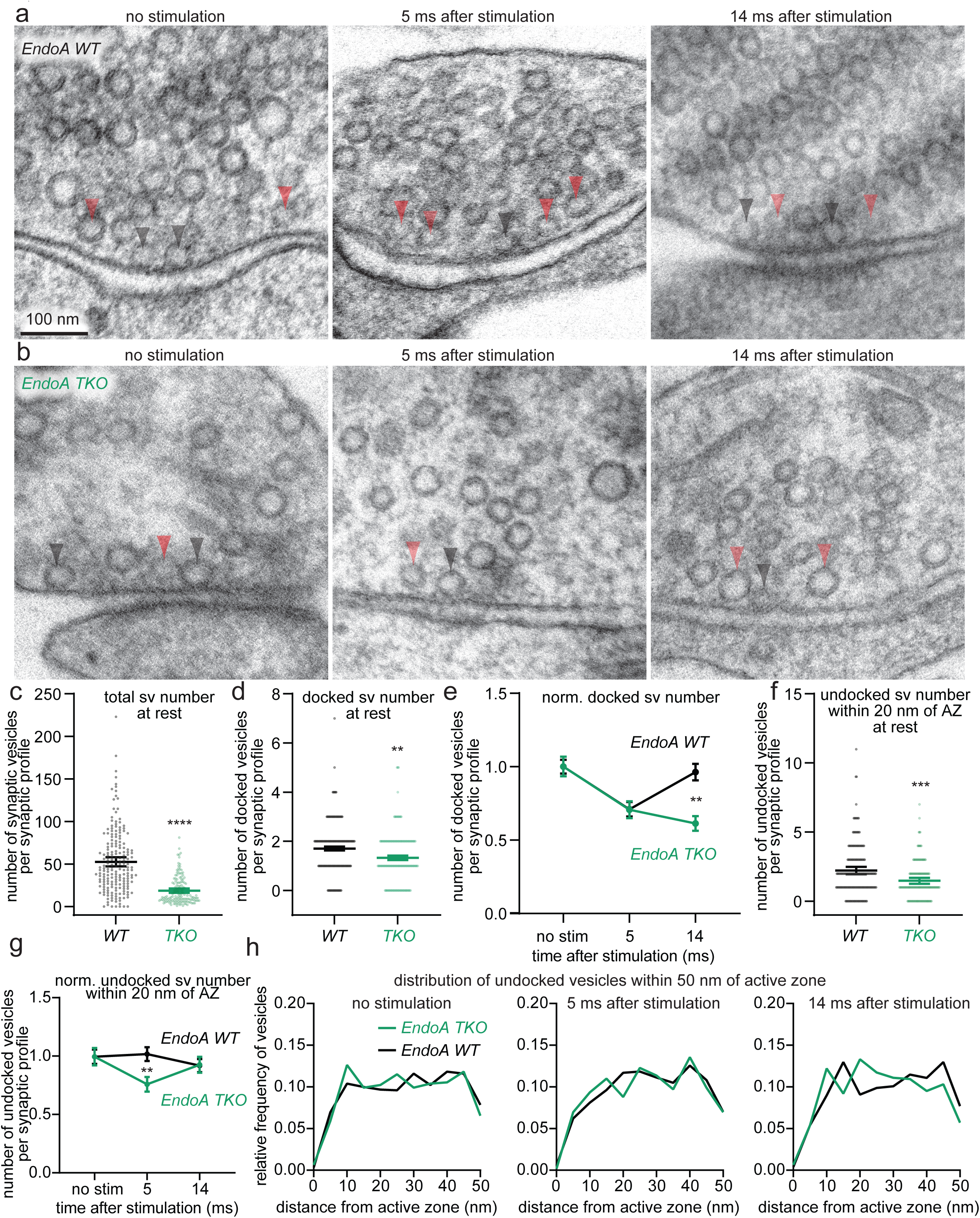
Endophilin A1 is required for transient docking of synaptic vesicles. a-b. Electron micrographs showing the progression of docked vesicle and undocked vesicle abundance and localization in *EndoA WT* (a) and *EndoA TKO* (b) synapses at rest, 5 ms, and 14 ms after stimulation (1 ms electric pulse, 37 °C, 1.2 mM external Ca^2+^). Black arrowhead: docked vesicle. White arrowhead: undocked vesicle. Scale bar, 100 nm. c-d. The total number of all vesicles in the terminal (c) and those docked in the active zone (d) at rest in *EndoA WT* (black) and *TKO* (green) synaptic profiles. Bars are the mean; error bars are SEM. Mann Whitney U test. ****p<0.0001, **p<0.01. e. Number of docked vesicle at rest, 5 and 14 ms after stimulation in *EndoA WT* and *EndoA TKO* synaptic profiles. 5 ms and 14 ms vesicle counts are normalized (norm.) to corresponding vesicle counts in the no stimulation (no stim.) control. Bars are the mean; error bars are SEM. Kruskal-Wallis test, with Dunn’s multiple comparisons test. *p<0.05, **p<0.01. Comparisons were made between *EndoA WT* and *EndoA TKO*. f. Number of undocked vesicles within 20 nm of the active zone membrane at rest in *EndoA WT* and *EndoA TKO* synaptic profiles. Bars are the mean; error bars are SEM. Mann-Whitney test. ***p<0.001. g. Number of undocked vesicles within 20 nm of the active zone at rest and 5, or 14 ms after stimulation in *EndoA WT* and *EndoA TKO* synaptic profiles. 5 ms and 14 ms vesicle counts are normalized to corresponding no stim vesicle counts. Bars are the mean; error bars are SEM. Kruskal-Wallis test, with Dunn’s multiple comparisons test. **p<0.01. Comparisons were made between *EndoA WT* and *EndoA TKO*. h. Relative frequency distributions of undocked vesicles 2-50 nm from the active zone membrane in *EndoA WT* and *EndoA TKO* synaptic profiles at rest, 5, and 14 ms after stimulation (left to right). Vesicle counts were separated into 2 nm bins and normalized by the total number of vesicles in this region. See Supplementary Table 1 for additional information.

### Endophilin A1 mobilizes replacement vesicle pool

Given its colocalization with Itsn1 in synapses and on vesicles, and the importance of the Itsn1-EndoA1 interaction in replacement vesicle localization and transient docking, we next assessed the contribution of EndoA directly to this replacement pathway. We compared Endophilin A (EndoA) triple knockout (TKO; *Sh3gl2-/-, Sh3gl1-/-, and Sh3gl3-/-*) mouse hippocampal neurons to WT neurons by zap-and-freeze. As in our previous studies^52^, the number of synaptic vesicles was reduced in EndoA TKO neurons by ∼70% due to the defect in synaptic vesicle recycling^52^ (Fig. 5a,b,c, and Extended Data Fig. 6a,b). Despite this strong reduction in total vesicles, the number of docked vesicles and replacement vesicles at rest was only modestly decreased (∼30%; Fig. 5a,b,d,f and Extended Data Fig. 6a,b,d), suggesting that remaining vesicles in EndoA TKO relatively accumulate near the active zone. Thus, vesicle counts were normalized to no stimulation conditions to account for baseline differences in resting synapses. The number of exocytic pits also decreased by ∼30% when compared to EndoA WT, but the frequency of their appearance still fell within the normal range^7^ (Extended Data Fig. 6c).

WT synapses displayed normal transient docking (Fig. 5a,e, and Extended Data Fig. 6a) and a subtle increase in replacement vesicles at 5 ms that returned to baseline at 14 ms (Fig. 5a,g, and Extended Data Fig. 6a), mirroring transient docking. Strikingly, in EndoA TKO synapses, transient docking completely failed (Fig. 5b,e, and Extended Data Fig. 6b). Further, following stimulation, undocked vesicles slightly decrease at 5 ms and return to baseline at 14 ms (Fig. 5b,g, Extended Data Fig. 6b), likely mirroring the failure of these vesicles to be used in transient docking. Concomitantly, the relative distribution of vesicles within the replacement zone in TKO neurons was similar to that in WT neurons, in both unstimulated synapses and those frozen 5 ms or 14 ms after stimulation (Fig. 5h). Together, these data suggest that replacement vesicles accumulate and are properly localized in EndoA TKO synapses but are not properly mobilized and thus, cannot be used for transient docking.

### Replacement vesicles are necessary for short-term plasticity and synaptic maintenance

To assess the importance of replacement vesicles for synaptic physiology, we measured field excitatory postsynaptic potentials (fEPSPs) in mouse hippocampal slices taken from Itsn1 WT and KO male or female littermates (P30-60) by stimulating the CA1 region and recording from CA1/CA2 synapses in the CA2 region (Fig. 6a). Our previous studies show two major physiological deficits in the absence of transient docking: no facilitation of synaptic transmission and faster synaptic depression^7,21^. These parameters can be assessed by measuring release probability and synaptic responses to paired stimuli (two stimulating pulses interspersed with various intervals) and to high-frequency stimulation (100 APs, 20 Hz). Consistent with our finding that the number of exocytic pits was unchanged in Itsn1 KO synapses (Fig. 3d), there was no difference in release probability between WT and Itsn1 KO slices (Fig. 6b). By contrast, Itsn1 KO showed only a slight reduction in paired-pulse response compared to WT (Fig. 6c), suggesting that synaptic transmission can facilitate despite the lack of transient docking, likely because the activity-dependent docking factor Syt7 is present in these synapses^21,53^. However, in response to a high-frequency train, facilitation early in the train (first 10 stimulations) was greatly reduced in Itsn1 KO slices (Fig. 6d-f, and Extended Data Fig. 7g), and consequently, this reduction led to a faster depression of synaptic signaling, suggesting that replacement vesicles are needed for synapses to keep up with instantaneous signaling demands. Later in the train (last 10 stimulations), the responses were more similar but still significantly different between WT and Itsn1 KO, with Itsn1 KO responses being smaller (Fig. 6d-f). Given that this phase represents the balance between exocytosis and replenishment, Itsn1 likely also plays a significant role in steady-state replenishment. In agreement, there was an appreciable reduction of overall release, synchronous and asynchronous release during the train, measured by charge transfer, in Itsn1 KO (Extended Data Fig. 7a-f). Despite these defects, synaptic recovery after the train was indistinguishable between WT and KO (Fig. 6g, and Extended Data Fig. 7h), suggesting Itsn1 KO synapses can steadily return to baseline release competency.

**Fig. 6.**
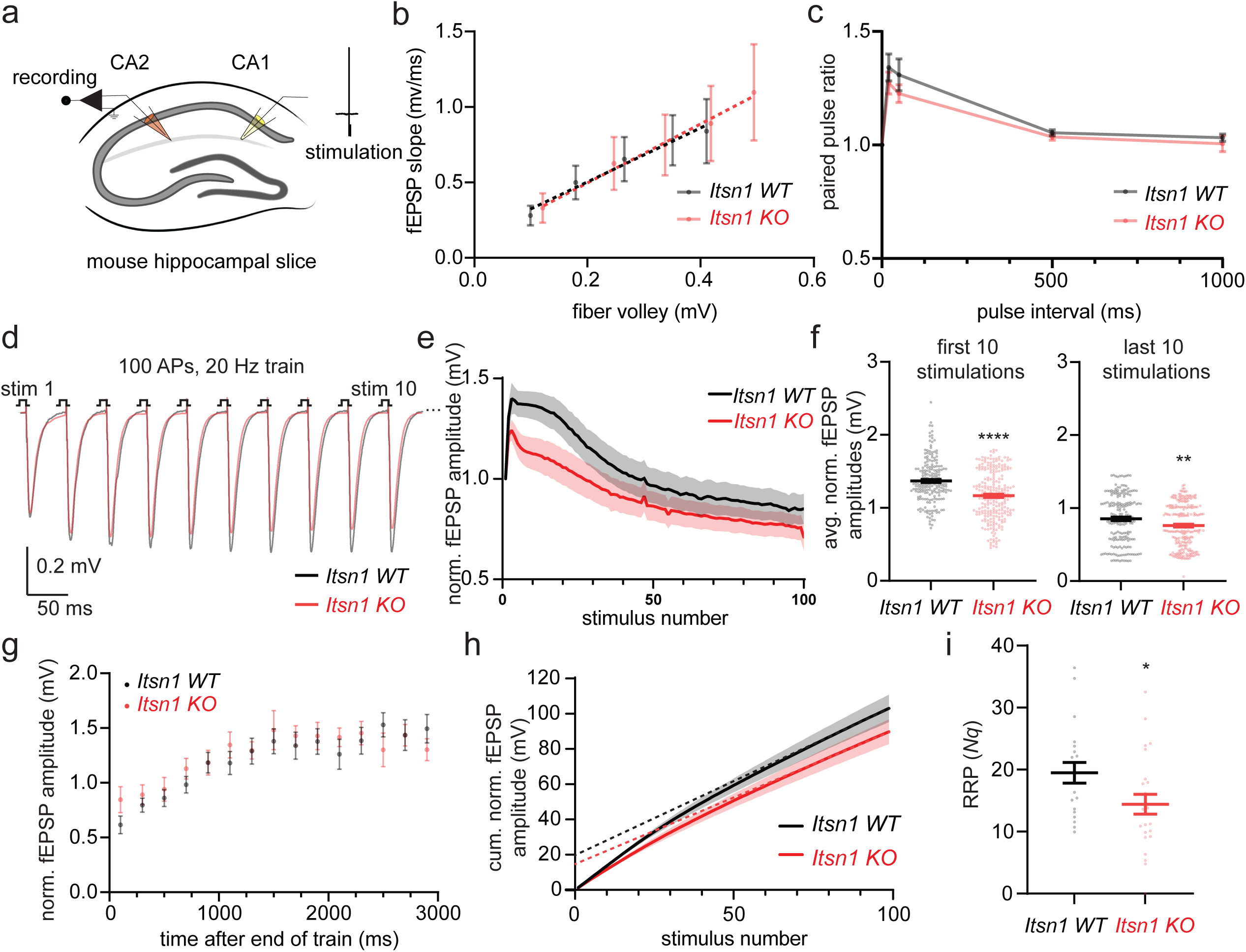
Itsn1 KO displays enhanced short-term depression and readily-releasable pool size reduction. a. Schematic of mouse hippocampal slice fEPSP recording procedure. A recording electrode was placed in the CA2/CA1 region of the hippocampus, and a stimulating electrode was placed in the CA1 region. 1 ms pulses were applied to slices and resulting field excitatory postsynaptic potentials (fEPSPs) were recorded. b. Release probability in *Itsn1 WT* and *Itsn1 KO* slices measured by comparing the fEPSP slope to fiber volley amplitudes in response to stimulations of increasing mV size (20 mV, 40 mV, 60 mV, 80 mV, and 100 mV). fEPSPs, field excitatory postsynaptic potentials. Dotted lines are linear regression analyses. Error bars are SEM. c. Plot showing paired-pulse ratio in *Itsn1 WT* and *Itsn1 KO* slices. A second pulse was separated by 20 ms, 50 ms, 500 ms and 1 s after the first pulse. The paired pulse ratio was determined by dividing the peak of the second pulse by the initial pulse peak. Dots are the mean; error bars are SEM. d. Average traces of the fEPSP response to the first 10 stimulations from a 100 action potential (AP), 20 Hz train applied to either *Itsn1 WT* (black) and *Itsn1 KO* (red) slices. *Itsn1 KO* responses began to decay by the 3^rd^ stimulation. e. Normalized (norm.) fEPSP response amplitudes through the course of the train stimulation in *Itsn1 WT* and *KO* slices. Transparent fill around traces is the SEM. f. The average (avg.) of normalized fEPSP amplitudes from the first 10 stimulations (left) and the last 10 stimulations (right) in *Itsn1 WT* and *KO* slices. Bars are the mean; error bars are SEM. Mann Whitney U test. ****p<0.0001. **p<0.01. g. Normalized fEPSP amplitude following 100 AP, 20 Hz trains in *Itsn1 WT* and *KO* slices, showing synaptic recovery. Recovery stimulations were initially applied 100 ms after the end of the train. Then, 200 ms were added sequentially to the time interval between the end of the train and the recovery stimulation out to 2900 ms. Dots are the mean, error bars are SEM. h. Cumulative (cum.) normalized fEPSP amplitudes during 100 AP, 20 Hz trains in *Itsn1 WT* and *KO* slices. Dotted lines are linear regression analyses. Transparent fill around traces is the SEM. i. Readily-releasable pool sizes (*Nq*) in synapses from *Itsn1 WT* and *KO* slices, approximated by linear back-extrapolation from traces in h. Mann Whitney test. *p<0.05. Bars are the mean; error bars are SEM. See Supplementary Table 1 for additional information.

Finally, we calculated the readily-releasable pool (RRP) size by linear back-extrapolation of cumulative normalized fEPSP responses during train stimulation^54–56^, analysis which relies on the assumption that synaptic depression coincides with RRP depletion. Thus, RRP can be taken as a reflection of the product of *N*, the number of release sites, and *q*, the quantal size (*Nq*) ^56–58^. Here, the RRP was reduced by 30% in Itsn1 KO slices when compared to WT slices (Fig. 6h,i). This reduction was consistent with a ∼35% reduction of replacement vesicles (Fig. 3h), suggesting that replacement vesicles make a substantial contribution to the RRP. Taken together, these data suggest that Itsn1 and EndoA1 maintain replacement vesicles within 20 nm of the active zone in the replacement zone, and these vesicles are used to replenish release sites during the train of stimuli to enhance synaptic signaling (Extended Data Fig. 7i).

## Discussion

### The interplay of synaptic phases

Phase separation is emerging as a major regulator of synaptic organization and function. Numerous presynaptic and postsynaptic proteins have been shown to phase separate *in vitro*, in heterologous cell systems, and in native contexts^30–32,34–36,40,59–61^. In the presynapse, many distinct compartments are thought to contain phase-separated proteins. In the active zone, proteins such as RIM, RIM-BP2, voltage-gated Ca^2+^ channels, Munc13, Liprin-ɑ, and ELKS-1 phase separate^30–33^, and this action has been linked to the development and maturation of the active zone, vesicle tethering, its organization, and structure. Adjacent to the active zone is the endocytic zone, which is recently shown to contain phase-separated Dynamin 1xA and Syndapin 1, critical for endocytic function^34^. Further into the synapse is the phase-separated synaptic vesicle reserve pool, which is largely achieved by the multivalent interactions of Syn1^35^. Here, we describe a zone driven by condensation of Itsn1 and EndoA1, positioned between the active zone and Syn1-labeled reserve pool cluster.

This tiered molecular organization seems to organize functional pools of synaptic vesicles. Vesicle pools within presynapses have been categorized into three main functional groups–the reserve pool, the recycling pool, and the readily-releasable pool^11,16,26,27^. Our data here suggests that the readily-releasable pool is further divided into the docked vesicle pool and the replacement pool (Extended Data Fig. 7i). Consistent with this idea, morphological studies in hippocampal mossy fiber synapses suggest that if all vesicles within 60 nm of the active zone were to fuse, the amount of membrane deposited into the presynaptic membrane would mirror the amount of membrane added during readily-releasable pool depleting stimulation, which was determined by capacitance measurements^25,62^. These molecular layers, and thereby functional pools, likely retain their identity in resting synapses due to the presence of physical boundaries (phase separations). Elevation of calcium during synaptic activity can potentially break these boundaries either directly by resolving the phase separation or indirectly by posttranslational modifications (as in CaMKII phosphorylation of Synapsin), enabling more vesicles to be available commensurate to the activity level of synapses. Thus, this tiered organization can modulate the number of available vesicles at a given synapse.

Nonetheless, how the size and durability of these phase-separated domains is controlled, and how they stay localized in synapses remains enigmatic. Recent work suggests that the size and stability of phase-separated domains can be determined via Pickering emulsions - interactions with immiscible protein clusters that localize to the surface of said domains^63,64^. Similarly, our results in HEK293 cells showed that Syn1-Itsn1 condensates containing FL Itsn1 were smaller in size when compared to Syn1-Itsn1 condensates that contained either fragment of Itsn1 (AB or AE). Likewise, in lamprey giant reticulospinal synapses, Itsn1 is found to increase the size of Syn1 condensates or disperse them, depending on abundance of Itsn1^36^. Thus, Itsn1 could regulate the size and stability of Syn1 condensates, and thereby the reserve pool vesicle cluster, similarly as described for several other phase-separating systems^64^. Such a function may directly relate to the size of the replacement pool described here. Further, interactions between these proteins may localize them to the presynapse. Itsn1 is long shown to interact with various transmembrane proteins in the active zone, like SNAP-25^45,65^. This may work to anchor Itsn1, and thus Syn1, along with their associated vesicle pools, at presynaptic terminals in a specific order from the active zone membrane.

### Crosstalk between functional vesicle pools

Three lines of evidence presented here suggest that the replacement zone is refilled from an upstream pool of vesicles, either the recycling or the reserve pool, in an activity-dependent manner, First, WT synapses typically display a transient increase in undocked vesicles within 20 nm of the active zone, without an overall decrease in undocked vesicle number within 50 nm, implying vesicle movement into this 20 nm region. Second, in Itsn1 KO synapses, replacement vesicles can eventually populate within 20 nm of the active zone, albeit on a timescale (∼100 ms after stimulation) incapable of accommodating synaptic demand for docked vesicles. This transition may be facilitated by the movement of new vesicles from further within the synapse into or near the replacement zone. Third, upon high-frequency stimulation, vesicles within the replacement zone are slightly reduced but largely maintained, implying the presence of an active refilling mechanism. These are consistent with previous optical studies^66,67^, and electrophysiological studies paired with mathematical modeling^20,22–24^. Taken together, our data suggest that the replacement zone consists of undocked vesicles, which can be used for the replacement of vacated release sites when the vesicles arrive within 20 nm of the active zone.

Currently, the upstream pool of vesicles is poorly defined. One possibility is that vesicles in the reserve pool can be accessed during activity. Consistent with this idea, Itsn1 and Syn1 interaction has been shown to be activity-dependent^46^, perturbing the Syn1-Syn1 interaction within the reserve pool and thereby, freeing vesicles for entry into the replacement pool and subsequent use in transmission. Here, we show that Itsn1 and EndoA1 localize at the periphery of the Syn1 reserve pool. Taken together with our finding that Itsn1 condensates are linked to vesicle clusters by Syn1, it is enticing to postulate that Itsn1-EndoA1 sit at the edge of the reserve pool, capturing periphery vesicles and using them for release site replenishment (Extended Data Fig. 7i). Alternatively, the recycling pool may provide vesicles to the replacement pool. Itsn1 and EndoA1 may potentially prevent the formation of Syn1-Syn1 interactions, which normally “lock” vesicles into the reserve pool^35,36,68,69^. Thus, vesicles that contain Itsn1 and EndoA1 after recycling are primed to reenter the readily-releasable pool. This transition may be further facilitated in glutamatergic neurons by the presence of vGlut1 on these vesicles, which interacts with both Itsn1 and EndoA1 to bolster evoked release capacity at synapses^70^. Thus, the Itsn1-EndoA1 replacement pool may contain vesicles poised to immediately refill vacated sites and upstream vesicles recently regenerated from synaptic vesicle recycling.

### Replacement vesicle mobilization

The persistence of replacement vesicles in EndoA TKO synapses even during activity sheds light on the mechanism of the replacement vesicle to docked vesicle transition. Our data suggests Itsn1 alone can fill the replacement pool at rest, but without EndoA, Itsn1 may “lock” these vesicles in the replacement pool, similarly to Syn1. In this model, Itsn1 generates the replacement pool, but hinders the usage of these vesicles, perhaps through phase separation of replacement vesicles (Extended Data Fig. 7i). EndoA may then through its interaction with Itsn1 free these vesicles from the replacement pool during activity for transient docking. Since EndoA1 interacts with vGlut1 to control release probability of vesicles, further dissecting how these molecular players interplay will heighten our understanding of neurotransmission and synaptic plasticity.

## Supporting information

Exnteded Data Figures

## Acknowledgements

We thank Dr. Volker Haucke for sharing the Itsn1 antibody. We also thank Dr. Melanie Pritchard from Monash University for sharing the Itsn1 KO animals and Dr. Sanda Predescu from Rush University for shipping the transgenic animals. We are also indebted to Dr. Johanna Pena Del Castillo for the help with endophilin dispersion experiments, and the past and current members of the Watanabe laboratory, particularly Dr. Kie Itoh, Sydney Brown, and Christian Pearson for technical assistance and Dr. Grant Kusick for discussion; M. Delanoy, B. Smith and Hoku West-Foyle at the Johns Hopkins Microscopy Facility, and Alexis Tomaszewski and Kristhine Martinez for the initial shRNA experiments. We thank the Advanced BioMedical Imaging facility at Charité for the support in microscopy and G. Schneider from the Milovanovic lab for the experimental support. S.W. and this work were supported by start-up funds from the Johns Hopkins University School of Medicine, Johns Hopkins Discovery funds, Johns Hopkins Catalyst award, Marine Biological Laboratory Whitman Fellowship, Chan-Zuckerberg Initiative Collaborative Pair Grant, Chan-Zuckerberg Initiative Supplement Award, Brain Research Foundation Scientific Innovation Award, Helis Foundation award, and the National Institutes of Health (1DP2 NS111133-01, 1R01 NS105810-01A1, R35 NS132153) awarded to S.W. S.W. is an Alfred P. Sloan fellow, a McKnight Foundation Scholar, a Klingenstein and Simons Foundation scholar, and a Vallee Foundation Scholar. T.H.O. was supported by the National Science Foundation GRFP (2019241734) and a grant from the National Institutes of Health to the BCMB program of the Johns Hopkins University School of Medicine (T32 GM007445). R.P. is supported by the American Heart Association Postdoctoral fellowship. A.H. was supported by the Albstein research scholarship. I.M. and S.G. received support from the Wellcome Trust (224361/Z/21/Z), John Fell (10447), FCT (PTDC/MED-NEU/8030/2020) and La Caxia (HR22-00854) Foundations and EU-Horizon 2020 (857524). B.H.C. funded/supported by the Deutsche Forschungsgemeinschaft (SFB1286/A01). D.M. is supported by the start-up funds from DZNE, the grants from the German Research Foundation (SFB 1286/B10 and MI 2104), and the Human Frontiers Science Organization (RGEC32/2023). C.H. is supported by a fellowship of the Innovative Minds Program of the German Dementia Association; HW is supported by the Oversea Study Program of Guangzhou Elite Project (SJ2020/2/JY202025). The EM ICE high-pressure freezer was purchased partly with funds from an equipment grant from the National Institutes of Health (S10RR026445) awarded to Scot C Kuo.

## Author Contributions

T.H.O., I.M., D.M., and S.W. conceived the study and designed the experiments. S.W. oversaw the overall projects and funded the research. I.M. and D.M. also provided funding. T.H.O., A.H., S.R., B.H.C. and S.W. performed electron microscopy experiments. T.H.O., C.H., W.W., A.H., and D.M., performed experiments concerning phase separation. T.H.O. performed superresolution imaging. S.G. and I.M. performed immuno-staining of purified vesicles and protein dispersion analysis. R.P. performed electrophysiology experiments. Co-second authors C.H., R.P., and H.W. contributed equally, and the order is simply alphabetical. Co-senior authors: I.M. and D.M. contributed equally, and the order is also alphabetical.

## Materials and Methods

### Animal use

All the animal work was performed according to the National Institutes of Health guidelines for animal research with approval from the Animal Care and Use Committees at the Johns Hopkins University School of Medicine. For Itsn1 KO experiments, 129SV/J^Itsn1-^ mice were maintained as heterozygotes, and neurons were cultured from homozygous null P0 pups, with homozygous WT littermates used as controls. Neurons were cultured from E18 embryos from C57BL/6J mice except for the *EndoA* TKO experiments.

For EndoA TKO, experiments complied with the national animal care guidelines and were approved by the University Medical Center Göttingen board for animal welfare and the animal welfare office of the state of Lower Saxony (LAVES). Constitutive knockout mice for endophilin A1, A2 and A3, as originally described^44^, were used in two separate breeding schemes: (i) endophilin A1-/--A2-/- (hereafter 1,2 DKO; lethal by P19-20) and littermate A1+/-A2+/- mice were obtained from breeding A1+/-A2+/- mice, and (ii) perinatally lethal endophilin A1-/--A2-/-A3-/- (hereafter TKO) and littermate A1-/-A2+/+A3-/- mice were obtained from breeding A1-/-A2+/-A3-/- mice. WT controls were obtained from breeding endophilin A1+/--A2+/-A3+/- (genetic background ∼80% C57BL/6J/∼20% SV129) mice or are obtained from the C57BL/6J line. Both male and female mice were used for all experiments.

### Primary neuron culture

Primary hippocampal neurons were isolated from either E18 embryos or P0 pups of both genders. The brains were taken from animals and hippocampi were dissected under a binocular microscope. Dissected hippocampi were collected in ice-cold dissecting media (1 x HBSS, 1 mM sodium pyruvate, 10 mM HEPES, pH7.2-7.5, 30 mM glucose, 1% penicillin-streptomycin) and later digested with papain (0.5 mg/ml) and DNase (0.01%) for 25 min at 37 °C. Cells were then further dissociated by trituration using fire-polished Pasteur pipettes.

For high-pressure freezing experiments neurons were plated onto 6-mm sapphire disks (Technotrade Inc) coated with poly-D-lysine (1 mg/ml) and collagen (0.6 mg/ml) with a pre-seeded astrocyte feeder layer on it. For this, cortices were harvested from E18/P0 animals, and astrocytes were isolated with a treatment of trypsin (0.05%) for 20 min at 37 °C, followed by trituration. Astrocytes were seeded in T-75 flasks containing DMEM supplemented with 10% Fetal Bovine Serum (FBS) and 0.2% penicillin-streptomycin. After 2 weeks, astrocytes were plated on sapphire disks (50K/well). After 1 week in culture, astrocytes were incubated with 5-Fluoro-2′-deoxyuridine (81 µM) and uridine (204 µM) for at least 2 hours to stop mitosis. Prior to the addition of hippocampal neurons, the medium was changed to Neurobasal-A (Gibco) supplemented with 2 mM GlutaMax, 2% B27 and 0.2% penicillin-streptomycin.

For high-pressure freezing experiments of *EndoA TKO* neurons and their controls, the following protocol was used. Astrocyte feeder cells were prepared as detailed in Pyott and Rosenmund (2002). Hippocampi from transgenic animals were dissected and incubated for 25 min in HBSS (Sigma) with 0.5% papain (Worthington) at 37 °C. After washing, neurons were triturated with fire-polished Pasteur pipettes, counted with a hemacytometer, and plated on astrocyte microislands (Bekkers and Stevens, 1991) in the plating medium [neurobasal medium (Invitrogen, Carlsbad, CA) supplemented with B-27 (Invitrogen), 17.3 mm HEPES, 1% GlutaMax-I (Invitrogen), 1% penicillin/streptomycin (Invitrogen), 25 μM β-mercaptoethanol, and 100 nM insulin (Heeroma et al., 2004)]. Medium was exchanged after about 12 h with neuronal medium. Neurons were cultured for 12–14 days before being used for experiments (only islands containing single neurons were examined). Dissociated cultures of primary cortical neurons have been prepared as previously described^44^.

For fluorescence imaging, dissociated hippocampal neurons were seeded onto 18-mm or 25-mm coverslips (Carolina Biologicals) coated with poly-L-lysine (1 mg/ml, Sigma) at a density of 25-40 × 10^3^ cells/cm^2^ in Neurobasal media (Gibco) supplemented with 2 mM GlutaMax, 2% B27, 5% FBS and 1% penicillin-streptomycin (NM5) at 37 °C in 5% CO_2_. The next day, the media was changed to Neurobasal media with 2 mM GlutaMax and 2% B27 (NM0), and neurons were maintained in this medium until use. Half of the media was refreshed every week or as needed.

For biochemical experiments, dissociated hippocampal neurons were seeded on poly-L-lysine (1mg/ml) coated plates or dishes with Neurobasal media supplemented with 2 mM GlutaMax, 2% B27, 5% FBS serum, and 1% penicillin-streptomycin, at a density of 1 × 10^5^ cells/cm^2^. The next day, the medium was changed to Neurobasal medium containing 2 mM GlutaMax and 2% B27 (NM0), and neurons were maintained in this medium. Half of the media was refreshed every week or as needed. For Itsn1 KO and WT littermate cultures, tail clips were obtained from live P0 pups and genotyped as previously described. Brain tissues were harvested from correct genotypes and hippocampal neurons were prepared as described above.

### Plasmids

For rescue experiments, rescue protein-coding sequences were recombined by In-fusion seamless cloning into a lentiviral expression vector containing 3x NLS-EGFP under the control of a human Synapsin promoter. Downstream of the 3x NLS-EGFP is a P2A sequence, which lies directly upstream of the destination for rescue protein-coding sequences. Plasmids containing either FL Itsn1 isoform or the same isoform containing W949E, Y965E (ΔEndoA1) mutations were generated.

For localization experiments, a plasmid containing the FL Itsn1 isoform N-terminally tagged to GFP (GFP-Itsn1) was purchased from addgene (#47395). For EndoA1, an EndoA1 protein-coding sequence which was a gift from the Michael A. Cousin lab was dropped into an pEGFP-N1 CMV mammalian expression vector (Clontech) by In-fusion cloning, resulting in Endo A1 C-terminally tagged with EGFP (EndoA1-GFP).

For 1,6-Hexanediol experiments in neurons, GFP-Itsn1 was used with co-expression markers mCherry-Syn1.

For heterologous cell system experiments, we used the plasmids described above along with mammalian expression vectors containing mCerulean-Endo A1, BFP-Syn1, GFP-Itsn1 AB, GFP-Itsn1 AE, untagged Syph, and Syph-emiRFP670, cloned and/or maintained in the Milovanovic lab.

### Lentivirus preparation

Lentivirus containing either Itsn1 WT or Itsn1ΔEndoA1 rescue constructs were prepared as described previously. Briefly, either rescue construct along with two helper DNA constructs (pHR-CMV8.2 deltaR (Addgene 8454) and pCMV-VSVG (Addgene 8455)) at a 4:3:2 molar ratio was transfected into HEK293T cells using polyethylene amine. Culture supernatant containing the virus was collected 3 days after transfection and 20-fold concentrated using Amicon Ultra 15 10K (Millipore) centrifugal filter. Aliquots were flash-frozen in liquid nitrogen and stored in -80 °C until use.

### Lentivirus infection

Neuron cultures prepared for lentiviral infection were grown until DIV (days in vitro) 7. Titered lentivirus was added to wells at an amount that leads to nearly 100% infection efficiency. Successful infection was assayed by the expression of NLS-EGFP which is contained within rescue constructs. Additionally, western blot analysis was used to assess the rescue construct protein of interest expression.

### Transient transfection

For transient expression of proteins, neurons were transfected at DIV 9-16 by Lipofectamine 2000 (Invitrogen) according to the manufacturer’s manual. Prior to transfection, half of the media from each well was taken out and mixed with fresh NM0 (see above) that was left to warm to 37 C° and equilibrate with CO_2_ in an incubator (recovery media). The rest of the media was aspirated and replaced with fresh NM0 for transfection. Plasmids were diluted in NeuroBasal Plus (Gibco) media so that 1-2 µg of DNA would be added to each well. Prior to the addition, DNA was mixed with a solution containing Lipofectamine 2000 such that there was a 1:1 to 1:4 ratio of µg DNA to µL Lipofectamine. This mixture was added to each well and incubated for 4 hours. Afterward, the transfection media was removed and replaced with recovery media previously prepared. After 16-20 hours, neurons were either used for pharmacological treatment or fixed for immunofluorescence.

Transient expression of proteins in HEK293T/HEK293 cells occurred at most one day prior to experiments. Cells were grown on 35-mm glass bottom dishes until 80% confluency. A solution of Lipofectamine 2000 and DNA (3 µL:1 µg) in 200 µL Optimem was prepared according to the manufacturer’s manual. 0.5 µg of DNA was used. The solution was mixed and allowed to sit for 20 mins prior to addition to cells. Cells were incubated for 12-24 hours after addition at 37 °C and 5% CO_2_ prior to imaging.

### Pharmacology

For 1,6-Hexanediol (Sigma) experiments in neurons, transfected neurons were treated immediately prior to imaging. Neurons plated on 25-mm coverslips were mounted onto a metal ring sample holder containing 3/4 of the final volume of cell culture media. Upon initiation of the imaging experiment, the remaining 1/4 cell culture media was added from a stock solution that contains 28% 1,6 Hexanediol to make a final concentration of 7% 1,6 Hexanediol.

For HEK293T cells, cells were plated onto 35-mm glass bottom petri dishes (Cellvis). The stock solution was 16%, for a final concentration of 4%. Dishes contained 3/4 of the final volume of cell culture media. Upon initiation of the imaging experiment, the remaining 1/4 cell culture media was added from a stock solution that contains 16% 1,6 Hexanediol to make a final concentration of 4% 1,6 Hexanediol.

For HEK293 cell experiments, pre-warmed 1,6-Hexanediol was diluted to final concentration of 3% in DMEM (culture media) and loaded onto cells plated on 35-mm glass bottom petri dishes (Cellvis, US).

### Immunofluorescence

For immunofluorescence, experiments were performed with DIV 14-16 hippocampal neurons. Culture media was removed from the wells and fixed with 37 °C 1X PBS containing 4% paraformaldehyde and 4% sucrose for 20 minutes at room temperature. After fixation, cells were washed three times with 1x PBS. Next, cells were permeabilized by 0.2% Triton X-100 diluted in 1X PBS for 8 minutes. After three washes with 1X PBS, cells were blocked by 1% BSA in 1X PBS for 1 hour. Then, coverslips were transferred to a humidified chamber and placed face down on a drop of primary antibody solution. Primary antibodies were diluted 1:500 to 1:250 in a 1% BSA 1X PBS and cells were incubated at 4°C overnight. GFP-Itsn1 and EndoA1-GFP were stained by an anti-GFP rabbit polyclonal antibody (MBL International). Endogenous Bassoon protein was stained by an anti-Bassoon mouse monoclonal (Synaptic Systems) antibody. Endogenous Rim protein was stained by an anti-RIM1 mouse monoclonal (Synaptic Systems) antibody. Endogenous Synapsin protein was stained by an anti-Synapsin 1/2 guinea pig polyclonal (Synaptic Systems) antibody. Endogenous Synaptobrevin 2 protein was stained by an anti-Synaptobrevin 2 mouse monoclonal (Synaptic Systems) antibody. Endogenous Endophilin A1 was stained by an anti-Endophilin 1 guinea pig polyclonal (Synaptic Systems) antibody. Intersectin-1 was stained by an anti-Intersectin-1 rabbit polyclonal (Gift from Volker Haucke) antibody. Next, coverslips were washed with 1X PBS three times. Secondary antibodies were diluted in 1X PBS containing 1% BSA. For superresolution 2D STED imaging, an anti-rabbit Atto647N (Rockland) secondary antibody was used at 1:120 dilution and an anti-mouse Alexa594 (Invitrogen) secondary antibody was used at 1:1000. For ISIM, a 1:500 dilution was used for Alexa488, Alexa568, or Alexa647 secondary antibodies. Secondary antibody incubation was performed in a humidified chamber as described previously for 1 hour at room temperature. Following three 1X PBS washes, cells were rinsed with Milli-Q water and mounted on a glass slide containing a drop of ProLong Diamond Antifade Mounting media (Thermo Fisher). Mounting Media was allowed to solidify for 24 hours at room temperature in the dark before proceeding to STED imaging.

### Stimulated emission depletion microscopy (STED)

All 2D STED images were obtained using a home-built two-color STED microscope^34^. A femtosecond laser beam with a repetition rate of 80 MHz from a Ti:Sapphire laser head (Mai Tai HP, Spectra-Physics) is split into two parts: one part is used to produce the excitation beam, which is coupled into a photonic crystal fiber (Newport) for wide-spectrum light generation and is further filtered by a frequency-tunable acoustic optical tunable filter (AA Opto-Electronic) for multi-color excitation. The other part of the laser pulse is temporally stretched to ∼300 ps (with two 15-cm long glass rods and a 100-m long polarization-maintaining single-mode fiber, OZ optics), collimated, expanded, and wave-front modulated with a vortex phase plate (VPP-1, RPC photonics) for hollow STED spot generation to de-excite the fluorophores at the periphery of the excitation focus, thus improving the lateral resolution. The STED beam is set at 765 nm with a power of 120 mW at the back focal plane of the objective lens (NA=1.4 HCX PL APO 100×, Leica), and the excitation wavelengths are set as 594 nm and 650 nm for imaging Alexa594 and Atto647N labeled targets, respectively. The fluorescent photons are detected by two avalanche photodiodes (SPCM-AQR-14-FC, Perkin Elmer). The images are obtained by scanning a piezo-controlled stage (Max311D, Thorlabs) controlled by the Imspector data acquisition program.

### Data analysis of 2D STED images

A custom MATLAB code package was used to analyze overexpressed GFP tagged Itsn1 and EndoA1 protein distribution relative to the active zone marked by Bassoon, and synaptic vesicle pools marked by Synaptobrevin 2 in 2D STED images^34^. First, STED images were blurred with a Gaussian filter with radius of 1.2 pixels to reduce the Poisson noise, and then deconvoluted twice using the built-in deconvblind function: the first point spread function (PSF) input is measured from nonspecific antibody signal in the STED images, and the second PSF input is chosen as the returned PSF from the first run of blind deconvolution^34^. Each time, 10 iterations are performed. Presynaptic boutons in each deconvoluted image were selected within 30×30 pixel (0.81 mm^2^) ROIs based on the varicosity shape and bassoon signal. The active zone or synaptic vesicle pool boundary was identified as the contour that represents half of the intensity of each local intensity peak in the Bassoon and Synaptobrevin 2 channel, respectively, and the Itsn1 or EndoA1 protein foci are picked as local maxima. The distances between the protein foci centers and the active zone or synaptic vesicle pool boundary are automatically calculated correspondingly. Itsn1 and EndoA1 protein foci continuous with the edge of ROIs, and the Bassoon or Synaptobrevin 2 signals outside of the transfected neurons were excluded from the analysis. For each condition, roughly 80-100 boutons (n) were quantified from 3 different cultures (N). The MATLAB scripts are available by request.

### Instant Structured Illumination Microscopy (ISIM) imaging

Neurons were imaged by the VT-iSIM from BioVision Technologies. A 100x objective lens (NA=1.4) was used. MetaMorph Imaging software (MDS Analytical Technologies) was used for acquisition. ∼90 µm × 60 µm x-y fields of view were captured, with an x-y pixel size of 0.0650 nm. Z-stacks were taken with 100 nm spacing between z-slices. A z-depth of 3 µm was used. Fluorophores were illuminated by 525 nm, 605 nm, and 700 nm lasers at consistent powers. Images were then deconvoluted by Microvolution software 2021.04, using a pinhole radius of 320 nm, a pinhole spacing of 2500 nm. Deconvoluted images were then analyzed by ImageJ.

### Airyscan imaging and data analysis

For Airyscan imaging, samples were imaged in Zeiss LSM880 (Carl Zeiss) in Airyscan mode. For 1,6 Hexanediol experiments in neurons, fluorescence was acquired using a 63x objective lens (NA = 0.55) at 488×488 pixel resolution and a pinhole size above the lower limit for Airyscan imaging, as computed by ZEN software. Neurons transfected with GFP-Itsn1 and mCherry-Syn1 on DIV 9 were imaged at DIV 14-16 prior to the addition of 1,6 Hexanediol. The field of view depended on the size of the neuron imaged. Full Z-stacks were acquired. Afterward, 1,6 Hexanediol was added to the imaging chamber, and immediately after neurons were imaged by full Z-stack imaging to assess the dispersion of molecular condensates. For analysis, images were background corrected by a rolling ball radius of 50 pixels. Then, a 1.0 sigma gaussian blur was applied. After, lines with a pixel width of 10 were drawn parallel to the orientation of axons across condensates in Fiji. Intensity values for both GFP-Itsn1 and mCherry-Syn1 were measured before and after 1,6 Hexanediol addition. Mean intensity and intensity variance were measured along axons and used to calculate the coefficient of variation (CV) by dividing the variance by the mean. CV values were normalized by the average CV value of axons measured in each condition, as reported previously^40^.

For 1,6 Hexanediol experiments in HEK293T cells, fluorescence was acquired using a 63x objective lens (NA = 0.55) at 1024×1024 pixel resolution with the following settings: pixel dwell 0.24 µs and pinhole size above the lower limit for Airyscan imaging, as computed by ZEN software. Cells were transfected with GFP-Itsn1 and imaged following successful expression. Cells were recorded at 2 Hz for 1 minute. At 30 s, 1,6 Hexanediol was added to measure the dispersion of molecular condensates. Condensate-condensate fusion, if recorded prior to 1,6 Hexanediol addition, was isolated and assessed separately.

Internal FRAP experiments were conducted in cells similarly prepared for 1,6 Hexanediol experiments. Fluorescence was acquired using a 63x objective lens (NA = 0.55) at 1024×1024 pixel resolution with the following settings: pixel Dwell 0.24 µs and pinhole size above the lower limit for Airyscan imaging, as computed by ZEN software. Cells were recorded with an exposure of 600 ms for 100 frames, for a total imaging time of ∼1 minute. After frame 3, Itsn1 condensates were bleached with a 488 laser initially at a diameter of 800 nm, then at 1600 nm. The recovery of fluorescence was measured throughout the course of the experiment. Intensity values were transformed into fractional recovery over time to the maximum intensity value after bleaching from the initial intensity value following bleaching at frame 4. To measure the Tau of fluorescence recovery, fractional recovery values were fit by a non-linear one-phase association function.

### Live-cell confocal imaging

Live-cell confocal imaging and the 1,6-Hexanediol assay in HEK293 cells were performed as described previously^71^. In short, HEK293 cells were plated onto 35-mm glass bottom petri dishes (Cellvis, US) and transfected with plasmids as indicated in the text using Lipofectamine 2000 (Thermo Fisher). Pre-warmed 1,6-Hexanediol was diluted to final concentration of 3% in DMEM (culture media) and loaded to cells. Imaging was performed on the Eclipse Ti Nikon Spinning Disk Confocal CSU-X, equipped with OkoLab Live-cells incubator (for control of temperature at 37C°, 5% CO2), 2 EM-CCD cameras (AndorR iXon 888-U3 ultra EM-CCD), Andor Revolution SD System (CSU-X), objectives PL APO 60x/1.4 NA oil immersion lens. Excitation wave lengths were: 405-nm for BFP, 488-nm for GFP; 561-nm for mCherry, 640-nm for emiRFP. All images were analyzed with ImageJ (NIH).

### Live imaging by spinning-disk confocal microscopy and data analysis

The dispersion of GFP-labeled Itsn1 and EndoA1-mRFP proteins was imaged using the custom-built spinning-disk confocal system described previously^42^. Neurons with low fluorescence (protein expression) and coverslips with lower transfection efficiency were preferred. Clustering and dispersion were monitored upon field stimulation of neurons at 37 °C with 300 action potential at 10 Hz in Tyrode buffer (119 mM NaCl, 5 mM KCl, 25 mM HEPES buffer, 2 mM CaCl_2_, 2 mM MgCl_2_, 6 g/L glucose, pH 7.4) containing 2-amino-5-phosphonovaleric acid (APV, 50 μM) and 10 μm 6-cyano-7-nitroquinoxaline-2,3-dione (CNQX, 10 μM) to block recurrent activity. Intensity changes from region of interest (marked around the centroid of fluorescence intensity in synaptic boutons and near boutons along the axon) in time series images were analyzed using imageJ software and represented as mean± SEM.

### High-pressure freezing

75K hippocampal neurons cultured on sapphire disks were frozen using a high-pressure freezer (EM ICE, Leica Microsystems). For functional assessment of Itsn1 KO, Itsn1 KO cells along with WT littermates were prepared. For rescue experiments, lentivirus was added to KO wells for expression of Itsn1 rescue constructs. For functional assessment of EndoA1 KO, EndoA TKO cells were plated with WT littermates. For some experiments, ferritin (2 mg/ml) was used as a fluid-phase marker and added to the cells for 5 minutes prior to freezing. Cells were frozen in a physiological saline solution (140 mM NaCl, 2.4 mM KCl, 10 mM HEPES, 10 mM glucose; pH adjusted to 7.3 with NaOH, 300 mOsm) containing NBQX (3 µM, Tocris) and bicuculine (30 µM; Tocris), which were added to block recurrent synaptic activity. CaCl_2_ and MgCl_2_ concentrations were adjusted as needed for experiments (mentioned in the results section). Zap-and-freeze experiments were performed as described earlier (Kusick et al, 2020). After freezing, samples were transferred under liquid nitrogen to an automated freeze substitution system at -90 °C (EM AFS2, Leica Microsystems). Using pre-cooled tweezers, samples were quickly transferred to anhydrous acetone at -90 °C. After disassembling the freezing apparatus, sapphire disks containing cells were quickly moved to cryo-baskets containing freeze substitution solutions and left inside EM AFS2. Freeze substitution was performed in solutions containing 1% glutaraldehyde and 0.1% tannic acid in anhydrous acetone (solution A) and then 2% osmium tetroxide in anhydrous acetone (solution B), which had been stored under liquid nitrogen and then moved to the AFS2 immediately before use. The freeze substitution program was as follows: −90 °C for 36 hrs in solution A, paused to swap to solution B after 6x 30 min washes in −90 °C acetone, 5 °C h^−1^ to −20 °C, 12 h at -20 °C, and 10 °C h^−1^ to 4 °C. Afterward, samples were removed from the freeze substitution chamber and warmed at room temperature by 4x 20 min washes with acetone before infiltration and embedding. For this latter protocol, all the steps were performed in universal sample containers (Leica Microsystems) and kept covered in aclar film to prevent any evaporation.

### Sample preparation for electron microscopy

Following freeze-substitution, and washing, a 100% epon araldite (epon 6.2 g; araldite 4.4 g; DDSA 12.2 g, and BDMA 0.8 ml) solution was prepared and diluted by acetone to get 30%, 70% and 90% solutions. Samples were infiltrated for at least two hours at room temperature sequentially in 30% and 70% epon-araldite. Samples were then transferred to caps of polyethylene BEEM capsules with 90 % epon araldite and incubated overnight at 4 °C. The next day, samples were transferred to new caps with fresh 100 % epon araldite, changed every 2 hours 3x, after which samples were cured at 60 °C for 48 hours.

After resin was cured, 40 nm sections were cut using an ultramicrotome (EM UC7, Leica microsystems) and collected on single-slot copper grids coated with 0.7 % pioloform. The sections were then stained with 2.5 % uranyl acetate in 50 % methanol and 50% water solution.

### Electron microscopy imaging and data analysis

Samples were imaged on a Hitachi 7600 TEM equipped with an AMT XR50 camera run on AMT Capture v6 (pixel size = 560 pm), at 80 kV on the 100,000x setting. Samples were blinded before imaging. Synapses were identified by a vesicle-filled presynaptic bouton and a postsynaptic density. Postsynaptic densities are often subtle in our samples, but synaptic clefts were also identifiable by 1) their characteristic width, 2) the opposed membranes following each other closely, and 3) vesicles near the presynaptic active zone. 120-130 micrographs per sample of anything that appeared to be a synapse were taken without close examination. All images were from different synapses.

EM image analysis was performed as previously described^4,5,7,21,34,52,72,73^. All images from a single experiment were randomized for analysis as a single pool. Only after this randomization were any images excluded from analysis, either because they appeared to not contain a bona fide synapse or the morphology was too poor for reliable annotation. The plasma membrane, the active zone, exocytic and endocytic pits, clathrin-coated pits, docked synaptic vesicles, and all synaptic vesicles in the bouton were annotated in ImageJ using SynapsEM plugins: [https://github.com/shigekiwatanabe/SynapsEM (copy archived at) <u>swh:1:rev:11a6227cd5951bf5e077cb9b3220553b506eadbe</u>]^73^. To minimize bias and error and to maintain consistency, all image segmentation, still in the form of randomized files, was thoroughly checked and edited by a second member of the lab. Features were then quantitated using the SynapsEM^73^ family of MATLAB (MathWorks) scripts (https://github.com/shigekiwatanabe/SynapsEM). Example electron micrographs shown were adjusted in brightness and contrast to different degrees (depending on the varying brightness and contrast of the raw images), rotated, and cropped in ImageJ before being imported into Adobe Illustrator.

## Biochemical Methods

### Isolation of synaptic vesicles and clathrin-coated vesicles

Synaptic vesicles and clathrin-coated vesicles were isolated as reported previously^49^. Equal protein concentration of synaptic and clathrin-coated vesicles were used for immunoblotting experiments.

### Western blot analysis

Standard SDS-PAGE blot was used to analyze total protein levels. An electrophoresis system (BIO-RAD) was used to perform the separation using custom-prepared gels (4-15%, pH 8.8), depending on the size of a protein to be analyzed by immunoblotting. After electrophoresis, the proteins were transferred onto a nitrocellulose membrane using the transfer system (BIO-RAD). The membranes were further blocked in 5% milk prepared in 1x Tris-buffered saline and 0.1% Tween 20 (TBS-T, blocking buffer) at room temperature for 1 h, and subsequently incubated with primary and secondary antibodies (diluted in the blocking buffer). The proteins were detected using the Odyssey infrared imaging system (Li-COR) and analyzed using Image Studio Lite (a software package from LI-COR Biosciences) and/or ImageJ (http://rsb.info.nih.gov/ij/index.html). Both software were used to compare the density (i.e. intensity) of bands on a digital image of the Western blot.

### Immunoprecipitation

For Itsn1-Endo A1 protein binding assessment, we homogenized whole mouse brains (∼P60) in a homogenization buffer containing 0.32 M sucrose, 10 mM HEPES at 7.4 pH, and cOmplete protease inhibitor (Roche). Homogenates were centrifuged and separated from supernatants. Lysis buffer containing 20 mM HEPES at pH 7.4, 50 mM KCl, 2 mM MgCl_2_, and 1% Triton X-100 was added to homogenates for 1 hr with regular trituration on ice. To assess Itsn1 rescue construct binding to Endo A1, Itsn1 KO mouse hippocampal cultures were grown on 10 cm dishes (Corning) and infected by lentivirus containing rescue cassettes on DIV 7. On DIV 14, cells were lysed with the above buffer and triturated for 30 min on ice. Infected cells were compared to WT neurons and non-infected KO neurons from littermates. Cell or whole-brain lysates were spun down at 4 °C, and the supernatant was collected. Lysates were then bound to Dynabeads (Thermo Fisher) per the manufacturer’s protocol containing an anti-Endophilin 1 guinea pig polyclonal antibody (Synaptic Systems). Proteins were eluted from beads by heating in 2x SDS sample buffer diluted in water from a 4x stock (2 mL 1 M Tris HCL, pH 6.8, 0.8g 10% SDS, 2.4 mL 2-ßMe, 4 mL Glycerol, MilliQ up to 10 mL, Bromophenol blue powder) at 70 °C for 10 min. Heated samples were then loaded into precast Tris Glycine 4-20% gradient gels (Bio-Rad) and run at 140 V for 45 minutes. After, proteins were transferred to methanol-activated PVDF membranes by a wet transfer system at 200 V for 90 minutes. Membranes were blocked for 30 minutes at room temperature in Intercept Blocking buffer (Li-COR), and then transferred into primary antibody solution (anti-Endophilin 1 (1:1000), anti-Itsn1 (1:1000, Millipore), anti-ß Actin (1:5000, Synaptic Systems) in Intercept Blocking buffer) overnight at 4 °C (shaking). Membranes were then washed 3x for 5 mins in TBS-T, and then transferred to a secondary antibody solution which contains IRDye secondary antibodies (Li-COR) diluted to 1:10,000 in Intercept Blocking Buffer for 1 hr at room temperature. Signal was detected using Li-COR Odyssey Clx and quantification was done by Image Studio Lite from Li-COR.

### Electrophysiology and data analysis

For Itsn1 KO experiments, adult Itsn1 WT and KO mice of both sexes ranging from 6-8 weeks of age were anaesthetized using a combination of Isoflurane inhalation and Avertin injection. Mice underwent cardiac perfusion using chilled sucrose solution (10 mM NaCl, 2.5 mM KCl,10 mM Glucose, 84 mM NaHCO_3_, 120 mM NaH_2_PO_4_, 195 mM Sucrose, 1 mM CaCl_2_, 2 mM MgCl_2_) saturated with oxygen 5% / carbon dioxide 95% (carbogen). Brain was rapidly dissected, and hippocampi removed. Hippocampi were then embedded in premade agarose molds and sliced at 400 µm using Leica VT1200S vibratome at a speed of 0.05 mm/s and amplitude of 1.0 mm. Slices were then transferred to artificial cerebrospinal fluid (ACSF; 119 mM NaCl, 2.5 mM KCl, 1.3 mM MgSO_4_, 2.5 mM CaCl_2_, 26 mM NaHCO_3_, 1 mM NaH_2_PO_4_, and 11 mM D-glucose (315 Osm, pH 7.4) heated using a water bath to 32 °C saturated with carbogen. Slices were recovered at this temperature for 15 min before being removed from the bath and recovered for 1 hour at room temperature.

Recordings were performed at 32 °C in ACSF. Glass pipettes containing silver chloride electrodes were used to both stimulate and record. The stimulating electrode was filled with ACSF and placed in CA1 while the recording electrode was filled with 1 M NaCl was placed to record from CA1/CA2 synapses in the CA2. To record paired pulse measurements a bipolar square pulse of 0.3 ms at 60 mV was applied followed by second pulse at varying intervals ranging from 20-1000 ms. To assess release probability, we acquired data from a range of stimulation strengths, starting with 20 mV and increasing in intervals of 20 mV to 100 mV, which reveals a linear relationship between fEPSP slope fiber volley amplitude. The slope and y-intercept of this line were used to identify relative differences in release probability. To assess overall release, we used depressing train stimulation a bipolar square pulse of 0.3 ms at 60 mV was applied for 100 pulses at 20 Hz. To assess synaptic recovery after train stimulation a single pulse was applied at varying intervals following the end of the train (100 pulses, 20 Hz) at various intervals ranging from 100 ms to 3 s. Recordings were taken using Multiclamp 700B and Digidata 1550B and Clampex v11.2 software. Stimulus was applied using A-M Systems Isolated Pulse Stimulator Model 2100. 1-3 slices per mouse were recorded. Traces were analyzed using a combination of Clampfit software v11.2 and custom MATLAB code.

### Statistical analysis

Detailed statistical information is collated in Supplementary Table 1.

Internal FRAP experiments yielded Tau measurements that were compared using a student’s t-test, as Tau values were normally distributed. An alpha of 0.05 was set for null hypothesis testing. Different HEK293T cell cultures (N) were imaged on separate days. Condensates that were assayed (n) were taken from multiple cells within a culture.

Molecular condensates (n) were quantified from 3-4 biological replicates per condition. Each biological replicate was a separate HEK293 culture (N). Multiple comparisons were made by either a Kruskal Wallis test or one-way ANOVA, depending on normality of distributions. For Kruskal Wallis tests, each condition was compared by Dunn’s multiple comparisons test. An alpha of 0.05 was set for null hypothesis testing. 2D STED images were acquired from 2-3 biological replicates per condition. Each replicate was a dissociated mouse hippocampal culture (N) taken from different mice on different days. For each N, roughly 30 presynaptic bouton regions of interest (ROIs) (n) were imaged from multiple transfected cells. ROIs from each replicate were pooled and quantified as previously described^34^. An alpha of 0.05 was set for null hypothesis testing. For Itsn1 and EndoA1 foci distance statistical analysis, pooled distance measurements from each condition were assessed for distribution normality. A full-pairwise Kruskal Wallis test was performed. Afterward, each condition was compared by Dunn’s multiple comparisons test.

Coefficient of variation (CV) analysis was performed from 3 biological replicates. Each replicate was a dissociated mouse hippocampal culture (N). Multiple axons were measured per replicate (n). An alpha of 0.05 was set for null hypothesis testing. Calculated CVs were compared by paired student’s t tests.

For electron microscopy data, measurements were taken from roughly 100 synaptic profile micrographs (n) per condition. Replicate high-pressure freezing experiments (N) were conducted with cultures taken from different mice on different days. Sample sizes for each replicate were inferred from previous flash-and-freeze experiments as opposed to power analysis. An alpha of 0.05 was set for null hypothesis testing. For count data sets such as these, non-normal, nonparametric distributions are assumed and typical. However, means are best to represent central tendency, and these data are binomially distributed. So, an ANOVA test with a Brown-Forsythe correction and Games-Howell post hoc was conducted. In the case of electron microscopy data sets with measurements of 0, Brown-Forsythe correction fails. Therefore, statistical comparison by a Kruskal-Wallis test followed by Dunn’s multiple comparisons was used instead. In cases where only two samples were compared, a student’s t-test with Brown-Forsythe correction and Games-Howell post hoc was conducted. Data sets that contained measurements that were 0 were instead compared by a Mann-Whitney test.

For electrophysiological data, fEPSP measurements were taken 3 times per experimental condition, per hippocampal slice (n). ∼2-5 slices were taken from each animal (N), ∼6 of which were measured per genotype. An alpha of 0.05 was set for null hypothesis testing. To compare the average amplitude of the 10 first and last stimulations, fEPSP amplitudes taken from 100 AP, 20 Hz trains in either Itsn1 WT or KO slices were pooled and compared by an unpaired student’s t-test. For linear back extrapolation analysis, normalized cumulative peak amplitudes were taken from 100 AP, 20 Hz trains conducted on either Itsn1 WT or KO slices and back extrapolated by linear regression analysis. y-intercepts were pooled and compared between Itsn1 WT and KO measurements by an unpaired student’s t-test.

### Code availability

Custom R, Matlab and Fiji scripts for electron microscopy analysis are available at https://github.com/shigekiwatanabe/SynapsEM.

### Data availability

Original images used in this work will be uploaded to Figshare. Data will be available upon request.

