## Supplementary material for "Intersectin and Endophilin condensates prime synaptic vesicles for release site replenishment": Exnteded Data Figures

Extended Data Fig. 1

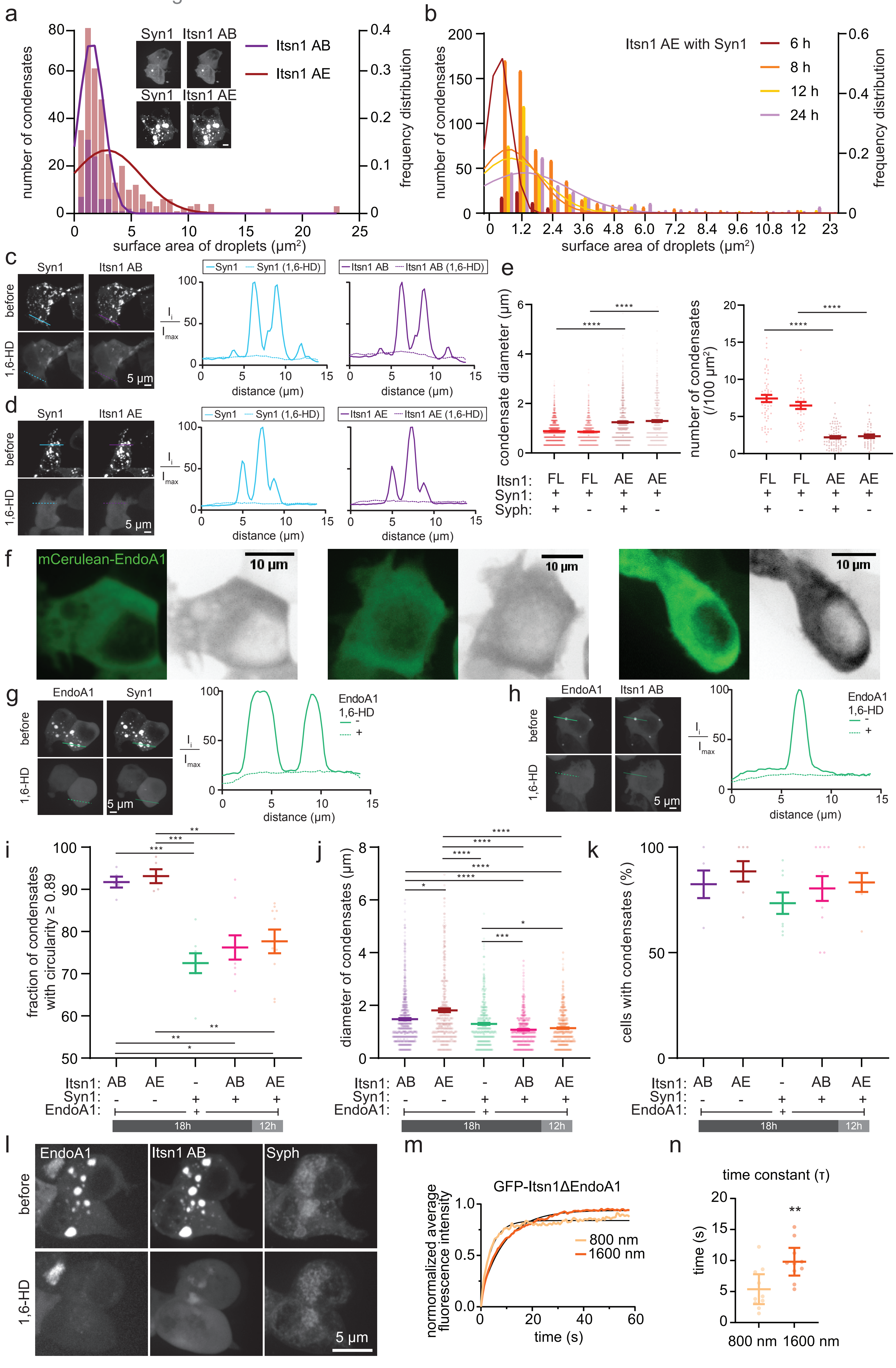

**Extended Data Fig. 1.**

a. A bar graph representing the number of condensates formed by co-expression of Syn1 with either a concatemer of two (Itsn1 AB) (purple) or five (Itsn1 AE) (red) Itsn1 Src-Homology 3 (SH3) domains, overlaid with the line graph representing the frequency distribution. Inset, representative images of HEK293 cells co-transfected with mCherry-Syn1 and GFP-Itsn1 AB or GFP-Itsn1 AE, respectively. Scale bar, 5 μm.

b. A bar graph representing the number of Itsn1 AE condensates with Syn1 (co-expression of GFP-Itsn1 and mCherry-Syn1) at the indicated time after transfection, overlaid with the line graph representing the frequency distribution. Note that the effective concentration increases with the time after transfection.

c. (left) Representative images of HEK293 cells co-expressing mCherry-Syn1 and GFP-Itsn1 AB domains before (top) and after application of 3% 1,6-Hexanediol (1,6-HD) (bottom). (right) plots showing fluorescence intensity of Syn1 (blue) and Itsn1 AB (purple) at the indicated line on the images before (solid line) and after (dotted line) the application of 1,6-Hexanediol. Scale bar, 5 μm.

d. Same as c, but with GFP-Itsn1 AE.

e. (left) Plot showing the condensate diameter of Itsn1 FL (red) /AE (dark red)-Syn1 condensates with or without co-expression of untagged Syph. Bars are the mean; error bars are SEM. Kruskal-Wallis test, with Dunn's multiple comparisons test. The condensates are analyzed from three independent transfections; p****<0.0001 was calculated when comparing Itsn1 FL-Syn1-Syph co-expressing cells to Itsn1 AE-Syn1-Syph co-expressing cells, and p****<0.0001 was calculated when comparing Itsn1 FL-Syn1 co-expressing cells to Itsn1 AE-Syn1 co-expressing cells. (right) Plot showing the number of Itsn1 FL/AE-Syn1 condensates with or without co-expression of untagged Syph. Bars are the mean; error bars are SEM. Kruskal-Wallis test, with Dunn's multiple comparisons test. The condensates are analyzed from three independent transfections; p****<0.0001 was calculated when comparing Itsn1 FL-Syn1-Syph co-expressing cells to Itsn1

AE-Syn1-Syph co-expressing cells, and p****<0.0001 was calculated when comparing Itsn1 FL-Syn1 co-expressing cells to Itsn1 AE-Syn1 co-expressing cells.

f. Three example images of live HEK293 cells expressing mCerulean-Endo A1. Scale bar, 10 μm.

g. (left) HEK293 cells co-expressing mCherry-Syn1 and mCerulean-EndoA1 before (top) and after application of 3% 1,6-Hexanediol (bottom). (right) plot showing fluorescence intensity of EndoA1 (green) at the indicated line on the images before (solid line) and after (dotted line) the application of 1,6-Hexanediol. Scale bar, 5 μm.

h. Same as g, but with GFP-Itsn1 AB and mCerulean-EndoA1.

i. Plot showing the circularity (≥ 0.89 ratio of x and y diameter, where x is the smaller axis) of either Itsn1-EndoA1, Syn1-EndoA1, or Itsn1-EndoA1-Syn1 condensates. The concatemer of Itsn1 was either AB or AE. Bars are the mean; error bars are SEM. Kruskal-Wallis test, with Dunn's multiple comparisons test. The condensates are analyzed from three independent transfections; p***<0.001 was calculated when comparing Itsn1 AB-EndoA1 to Syn1-EndoA1, p**<0.01 was calculated when comparing Itsn1 AB-EndoA1 to Itsn1 AB-Syn1-EndoA1, and p*<0.05 was calculated when comparing Itsn1 AB-EndoA1 to Itsn1 AE-Syn1-EndoA1. p***<0.001 was calculated when comparing Itsn1 AE-EndoA1 to Syn1-EndoA1, and p**<0.01 was calculated when comparing Itsn1 AE-EndoA1 to either Itsn1 AB-Syn1-EndoA1 or Itsn1 AE-Syn1-EndoA1.

j. Plot showing the diameter of condensates containing either Itsn1-EndoA1, Syn1-EndoA1, or Itsn1-EndoA1-Syn1. The concatemer of Itsn1 was either AB or AE. Bars are the mean; error bars are SEM. Kruskal-Wallis test, with Dunn's multiple comparisons test. The condensates are analyzed from three independent transfections; p*<0.05 was calculated when comparing Itsn1 AB-EndoA1 to Itsn1 AE-EndoA1, and p****<0.0001 was calculated when comparing Itsn1 AB-EndoA1 to either Itsn1 AB-Syn1-EndoA1 or Itsn1 AE-Syn1-EndoA1. p****<0.0001 was calculated when comparing Itsn1 AE-EndoA1 to either Syn1-EndoA1, Itsn1 AB-Syn1-EndoA1, or Itsn1 AE-

Syn1-EndoA1. p***<0.001 was calculated when comparing Syn1-EndoA1 to Itsn1 AB-Syn1-EndoA1, and p*<0.05 was calculated when comparing Syn1-EndoA1 to Itsn1 AE-Syn1-EndoA1.

k. Plot showing the fraction of cells containing either Itsn1-EndoA1, Syn1-EndoA1, or Itsn1-EndoA1-Syn1 condensates. The concatemer of Itsn1 was either AB or AE. Bars are the mean; error bars are SEM.

l. (top) HEK293 cells expressing mCerulean-EndoA1, GFP-Itsn1 AB, and Syph-emiRFP670. (bottom) cells in (top) after 3% 1,6 Hexanediol treatment. Scale bar, 5 μm.

m. Same as Fig.1f, but with GFP- Itsn1ΔEndoA1.

n. Same as Fig. 1g, but with GFP-Itsn1ΔEndoA1. Bars are the mean; error bars are SEM. Student's t test. p value **<0.01.

See Supplementary Table 1 for additional information.

Extended Data Fig. 2

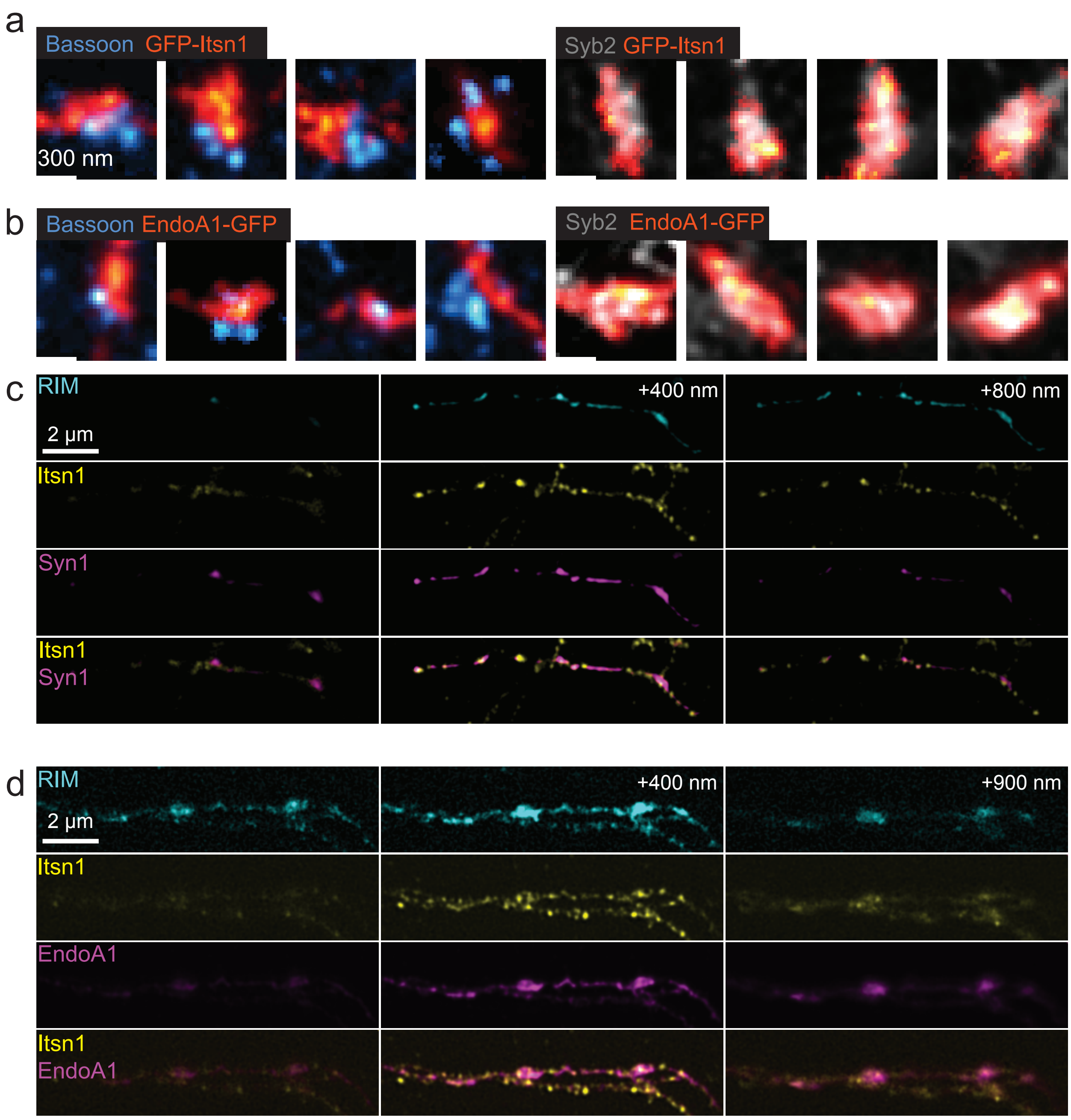

**Extended Data Fig. 2.**

a. Example 2D STED images of different mouse hippocampal presynapses expressing GFP-Itsn1 showing its localization relative to the active zone, marked by Bassoon (cyan) or synaptic vesicles, marked by Syb2 (gray). GFP-Itsn1 was stained by an anti-GFP antibody and a secondary antibody conjugated to Atto647N. Endogenous Bassoon was used as a proxy for the active zone and was stained by an anti-Bassoon antibody and a secondary antibody conjugated to Alexa594. Endogenous Syb2 was stained by an anti-Syb2 antibody and a secondary antibody conjugated to Alexa594. Scale bar, 300 nm.

b. Example 2D STED images of different mouse hippocampal presynapses expressing EndoA1-GFP showing its localization relative to the active zone, marked by Bassoon (cyan) or synaptic vesicles, marked by Syb2 (gray). EndoA1-GFP was stained by an anti-GFP antibody and a secondary antibody conjugated to Atto647N. Endogenous Bassoon was used as a proxy for the active zone and was stained by an anti-Bassoon antibody and a secondary antibody conjugated to Alexa594. Endogenous Syb2 was stained by an anti-Syb2 antibody and a secondary antibody conjugated to Alexa594. Scale bar, 300 nm.

c. Additional representative ISIM images showing RIM, Itsn1, and Synapsin 1, as in Fig. 2d. Side-view shown. Scale bar, 2 μm.

d. Additional representative ISIM images showing RIM, Itsn1, and EndoA1, as in Fig. 2f. En face view shown. Scale bar, 2 μm.

Extended Data Fig. 3

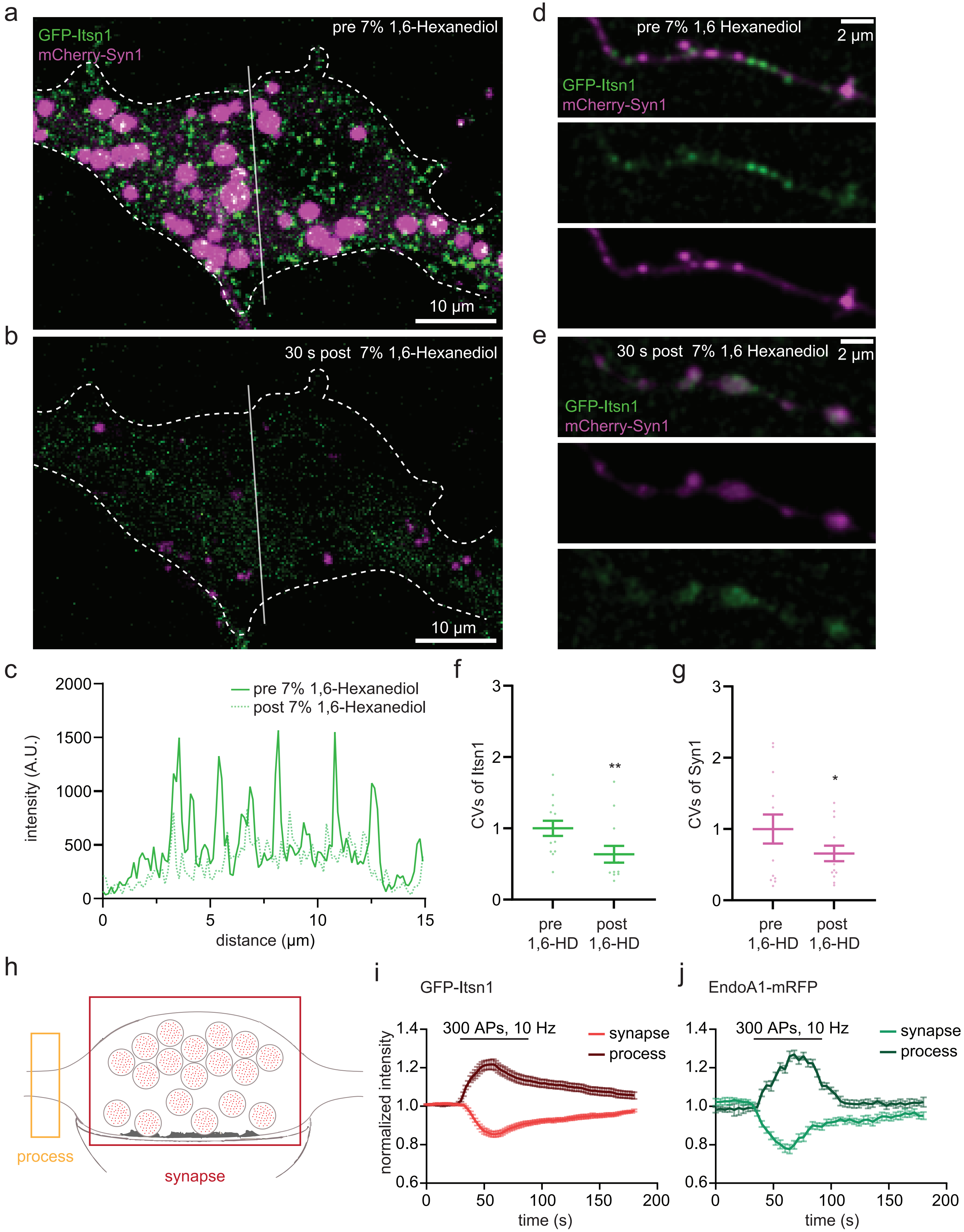

**Extended Data Fig. 3.**

a. Example micrograph of a neuron expressing GFP-Itsn1 mCherry-Syn1 before the application of 7% 1,6-Hexanediol. Dotted lines indicate cell boundary. Scale bar, 10 μm.

b. the same cell as in a, but 30 s after the application of 7% 1,6-Hexanediol. Dotted lines indicate cell boundary. Scale bar, 10 μm.

c. Line plots of fluorescence intensity from the transparent white line in (a) and (b) indicated before (green solid) and after (green dotted) the application of 7% 1,6-Hexanediol.

d. Example micrographs of an axon from a neuron expressing GFP-Itsn1 and mCherry-Syn1 before the application of 7% 1,6-Hexanediol. Scale bar, 2 μm.

e. The same axon in d, 30 s after the application of 7% 1,6-Hexanediol. Scale bar, 2 μm.

f-g. Coefficient of variation (CV) of GFP-Itsn1 (f) and mCherry-Syn1 (g).

h. Axonal (orange) and synaptic (red) regions used for fluorescence intensity measurements.

i. Plots showing GFP-Itsn1 fluorescence intensity at synaptic boutons or axonal processes. Fluorescence intensity was measured as depicted in (h) and normalized to the baseline. Traces shown are average values; error bars are SEM.

j. Plots showing EndoA1-mRFP fluorescence intensity at synaptic boutons or axonal processes. Fluorescence intensity was measured as depicted in (h) and normalized to the baseline. Traces shown are average values; error bars are SEM.

See Supplementary Table 1 for additional information.

Extended Data Fig. 4

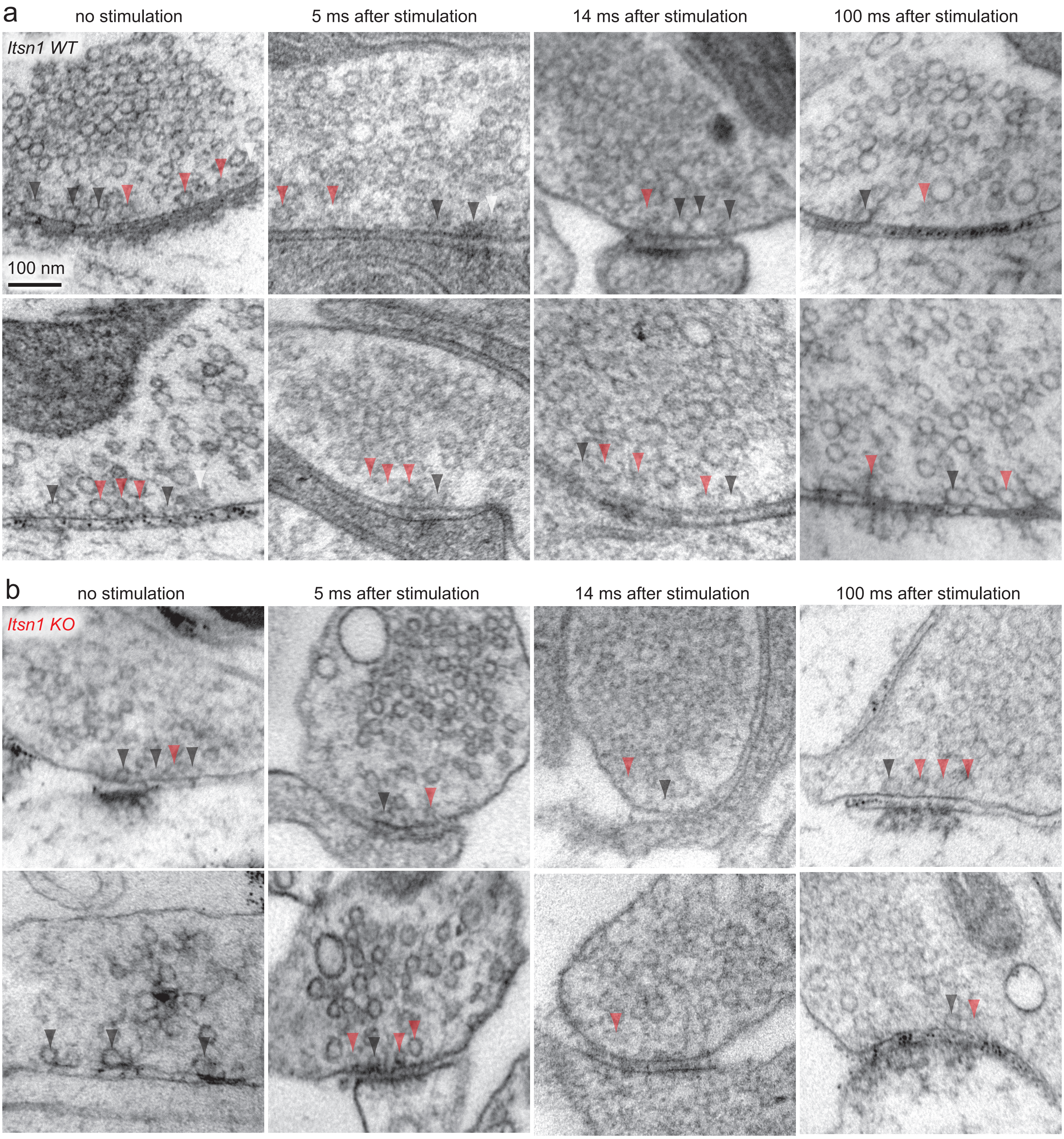

**Extended Data Fig. 4.**

a. Additional example electron micrographs from Fig. 3a. Scale bar, 100 nm.

b. Additional example electron micrographs from Fig. 3b.

Extended Data Fig. 5

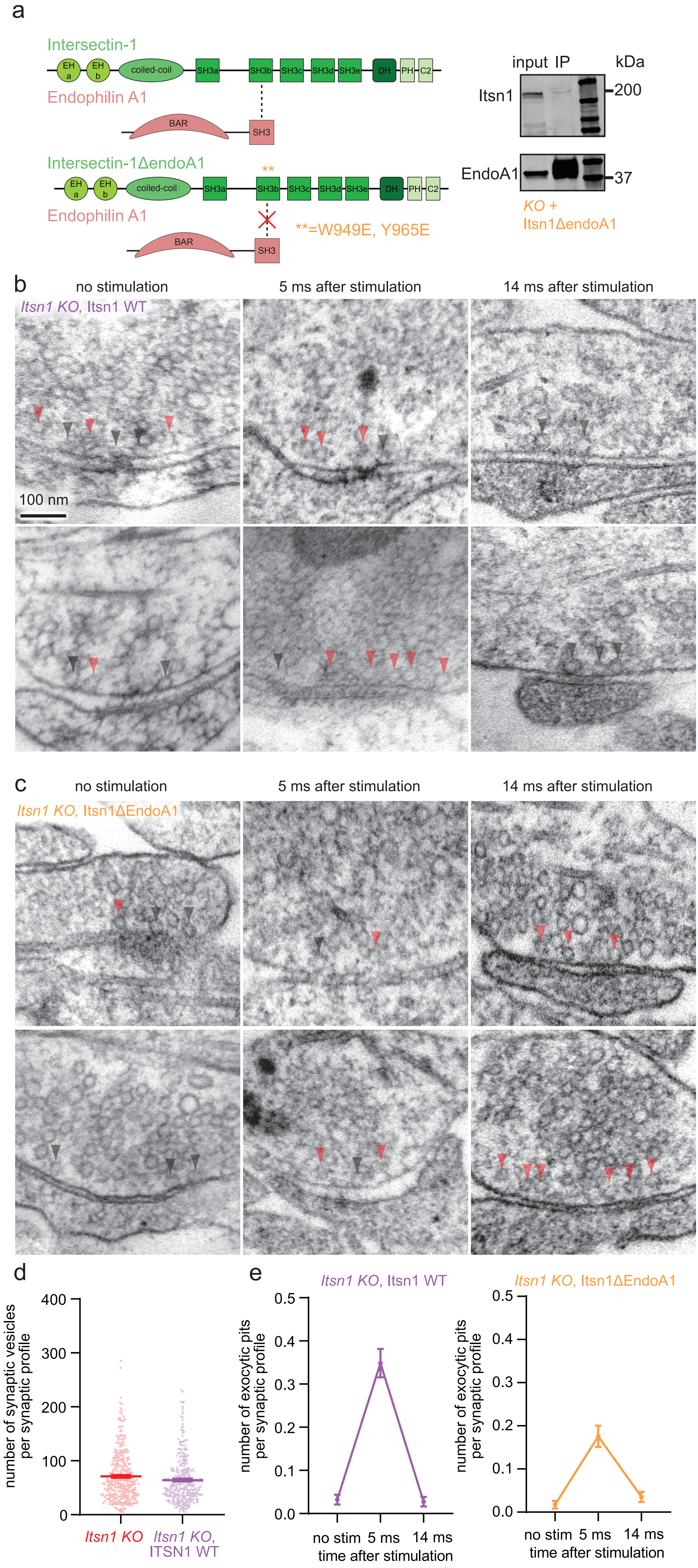

**Extended Data Fig. 5.**

a. Schematic showing Itsn1 and EndoA1 domain structures with the Itsn1-EndoA1 binding site denoted between participating domains by a dotted line, and Western blot showing the reduction of EndoA1 binding by Itsn1ΔEndoA1 mutations.

b. Additional example electron micrographs from Fig. 4a. Scale bar, 100 nm.

c. Additional example electron micrographs from Fig. 4b.

d. Number of synaptic vesicles per synaptic profile in *Itsn1 KO* (red) and *Itsn1 KO*, Itsn1 WT (purple) neurons.

e. Plots showing the number of exocytic pits in *Itsn1 KO*, Itsn1 WT (purple, left) and *Itsn1 KO*, Itsn1ΔEndoA1 (orange, right) synaptic profiles at rest, 5 ms, and 14 ms after stimulation.

Extended Data Fig. 6

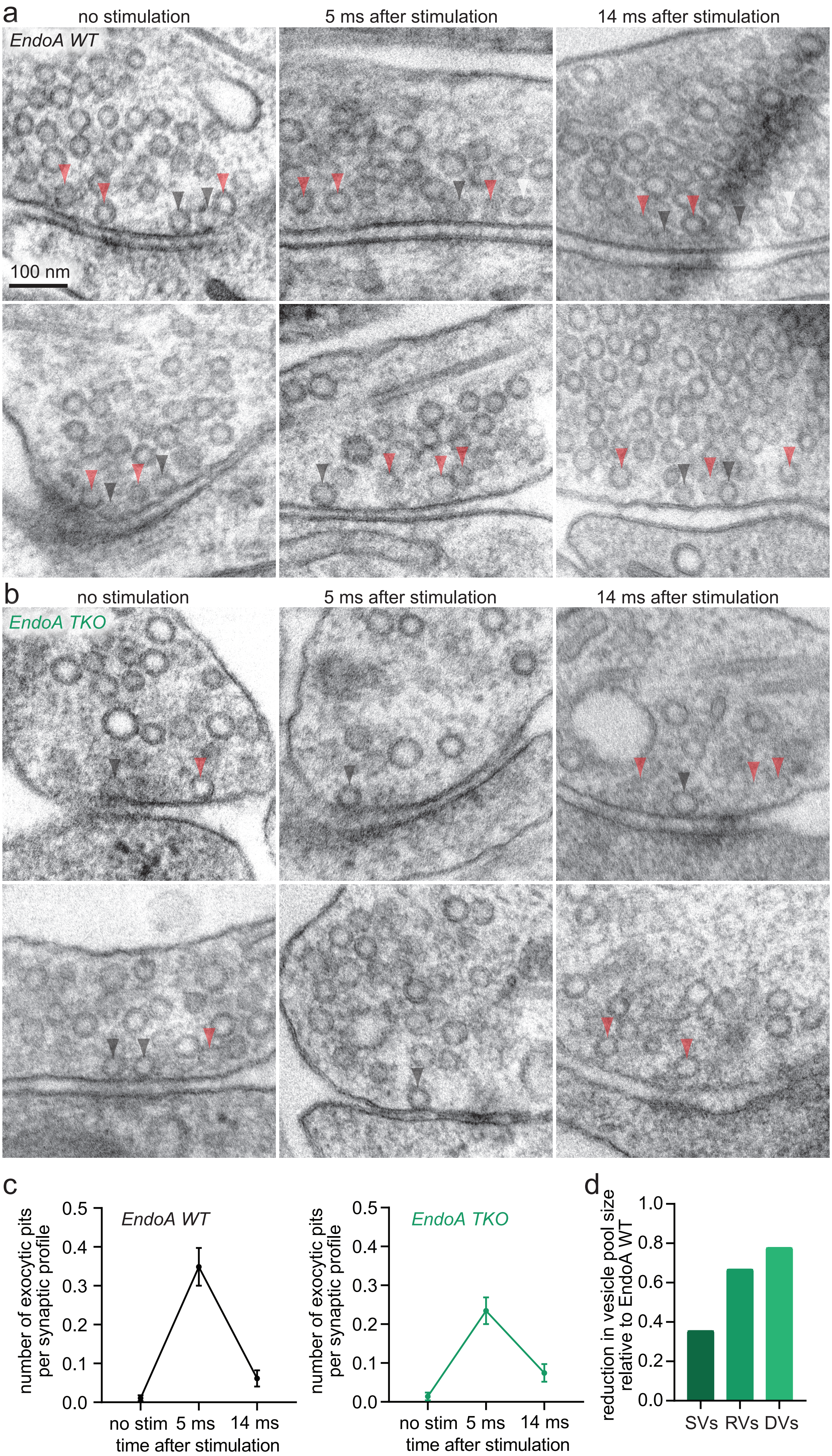

**Extended Data Fig. 6.**

a. Additional example electron micrographs from Fig. 5a. Scale bar, 100 nm.

b. Additional example electron micrographs from Fig. 5b.

c. Plots showing the number of exocytic pits in *EndoA1 WT* (black, left) and *TKO* (green, right) synaptic profiles at rest, 5 ms, and 14 ms after stimulation.

d. Bar graph showing the reduction in indicated vesicle pool sizes when calculating the ratio of vesicle counts between *EndoA TKO* and *WT*. SVs = synaptic vesicles, RVs = replacement vesicles, DVs = docked vesicles.

Extended Data Fig. 7

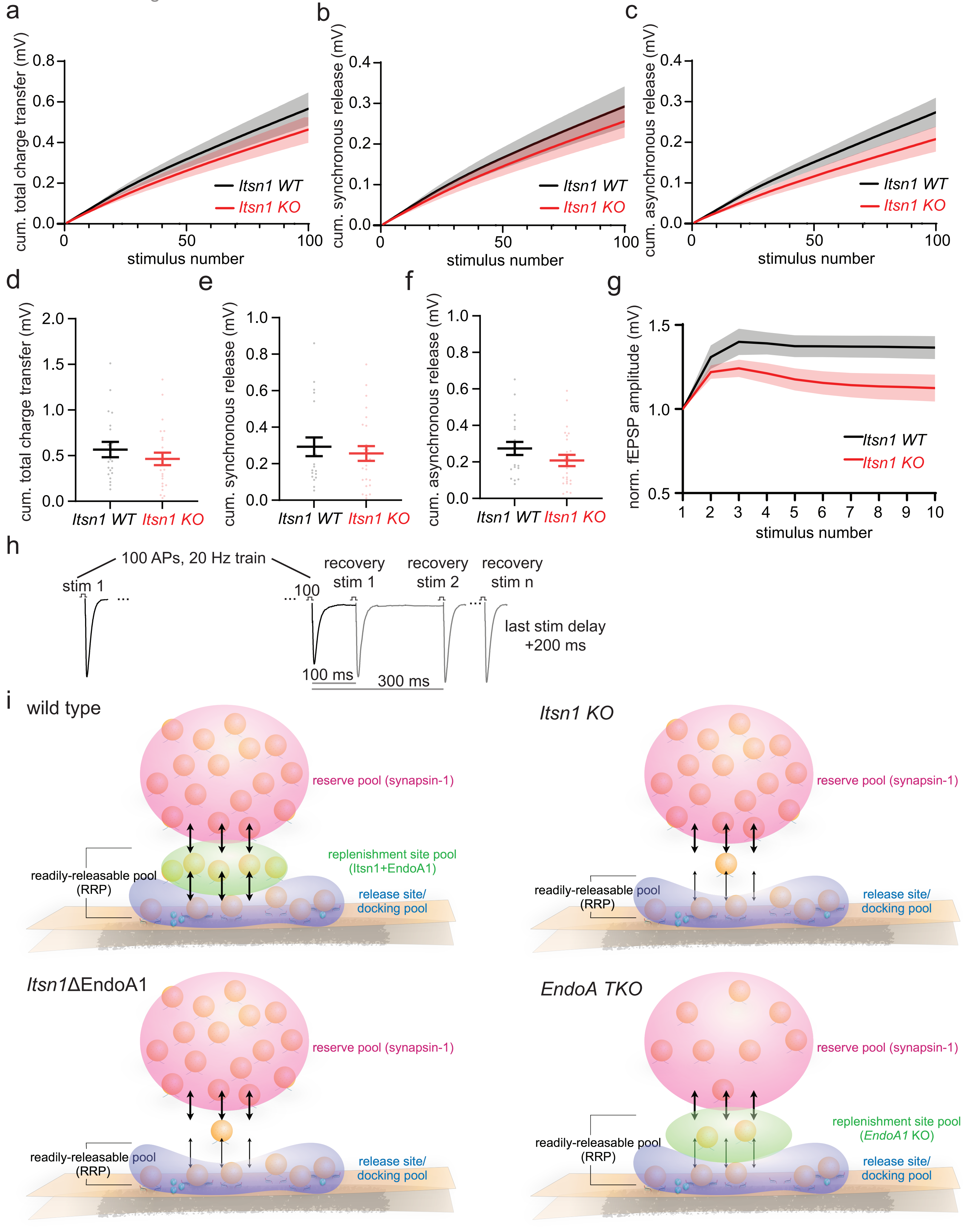

**Extended Data Fig. 7.**

a. Cumulative (cum.) total charge transfer (mV) measured from field excitatory postsynaptic potentials (fEPSPs) in response to stimulation; 100 AP, 20 Hz train. Transparent fill around traces is the SEM.

b-c. Cum. total charge transfer from a, separated by the first 10 ms from each pulse (b) and the next 40 ms (c). The first 10 ms is defined as synchronous, while the next 40 ms asynchronous. Transparent fill around traces is the SEM.

d. Last cum. total charge transfer values measured from fEPSPs in response to a 100 AP, 20 Hz train. Bars are the mean, error bars are SEM.

e. Last cum. synchronous (first 10 ms of release per stimulation) charge transfer values measured from fEPSPs in response to a 100 AP, 20 Hz train. Bars are the mean, error bars are SEM.

f. Last cum. asynchronous (last 40 ms of release per stimulation) charge transfer values measured from fEPSPs in response to a 100 AP, 20 Hz train. Bars are the mean, error bars are SEM.

g. Normalized fEPSP amplitude from the first 10 pulses during the train. Transparent fill around traces is the SEM.

h. Experimental paradigm schematic of synaptic recovery experiments in Fig. 6g. Following 100 AP, 20 Hz trains, single recovery stimulations were applied first at 100 ms after the train, then 300 ms, and then with delays increasing by 200 ms every next recovery stimulation n.

i. Schematic showing a new model of activity-dependent replenishment of release sites during short-term plasticity. The readily-releasable pool (RRP) consists of the docked vesicle pool and replacement vesicle pool. The replacement vesicle pool is maintained by Itsn1 and EndoA1, while the reserve pool is maintained by Synapsin 1. Following stimulation, the vacated release sites are rapidly replenished by replacement vesicles to enhance synaptic signaling. In the absence of Itsn1, the replacement pool is depleted. With only Itsn1ΔEndoA1 present, the replacement pool is also depleted. In *EndoA TKO*, despite a severe loss of vesicles in the terminal, the replacement pool is relatively maintained, but vesicles are not mobilized.
